# Molecular view of ER membrane remodeling by the Sec61/TRAP translocon

**DOI:** 10.1101/2022.09.30.510141

**Authors:** Sudeep Karki, Matti Javanainen, Dale Tranter, Shahid Rehan, Juha T. Huiskonen, Lotta Happonen, Ville O. Paavilainen

## Abstract

Protein translocation across the endoplasmic reticulum (ER) membrane is an essential initial step in protein entry into the secretory pathway. The conserved Sec61 protein translocon facilitates polypeptide translocation and coordinates cotranslational polypeptide processing events. In cells, the majority of Sec61 is stably associated with a heterotetrameric membrane protein complex, the translocon associated protein complex (TRAP), yet the mechanism by which TRAP assists in polypeptide translocation or cotranslational modifications such as N-glycosylation remains unknown. Here, we demonstrate the structure of the core Sec61/TRAP complex bound to a mammalian ribosome by Cryo-EM. The interactions with ribosome anchor the Sec61/TRAP complex in a conformation that renders the ER membrane locally thinner by significantly curving its the lumenal leaflet. We propose a model for how TRAP stabilizes the ribosome exit tunnel to assist nascent polypeptide insertion through Sec61 and provides a ratcheting mechanism into the ER lumen by direct polypeptide interactions.

## INTRODUCTION

Up to a third of eukaryotic proteomes are synthesized at the surface of the endoplasmic reticulum (ER) where they are initially inserted into the protein secretory pathway (Hegde and Keenan 2022); (Pool 2022). Most eukaryotic secretory proteins are targeted to the ER cotranslationally through recognition of N-terminal hydrophobic signal peptides or transmembrane segments by the signal recognition particle (SRP), which directs the ribosome nascent chain complex (RNC) for final delivery to the Sec61 protein translocon. SRP recognition of secretory proteins is based upon interaction with a nascent N-terminal cleavable hydrophobic signal peptide or transmembrane segment. The large sequence diversity of targeting peptides suggests that their targeting and insertion mechanisms may vary (Lang et al. 2022); (Liaci and Förster 2021)

The evolutionarily conserved heterotrimeric Sec61 channel is responsible for ER translocation of nascent polypeptides and is alone sufficient for translocation of nascent polypeptides across the ER membrane (Görlich and Rapoport 1993). However, a subset of secretory proteins contain signal peptides that are inefficient in engaging with Sec61 and cannot be translocated by Sec61 alone (Fons et al. 2003); (Hegde et al. 1998). Structural information about different states of the isolated heterotetrameric Sec61 translocon exist (Gemmer and Förster 2020), but several additional proteins can form either stable or dynamic complexes together with the core Sec61 complex, and presumably promote translocation of specific nascent proteins or to guide other processing events such as correct post-translational modifications during the critical membrane translocation process (Gemmer and Förster 2020). However, because of the difficulty inherent in isolating and characterizing higher order Sec61 complexes, structural information about their organization currently remains limited.

One complex that promotes Sec61-mediated insertion in many higher eukaryotes is the heterotetrameric translocon-associated protein (TRAP) complex (Fons et al. 2003); (Görlich and Rapoport 1993). Unlike most Sec61-associating protein, the TRAP complex forms a stable interaction with Sec61 (Hartmann et al. 1993); (Ménétret et al. 2008), and appears to form constitutive component of the Sec61 complex in cells (Pfeffer et al. 2017); (Braunger et al. 2018). Biochemical experiments suggested that TRAP may aid ER insertion of proteins with specific signal peptides (Nguyen et al. 2018) and modulate topogenesis of certain integral membrane proteins (Sommer et al. 2013).

In mammalian cells a subset of Sec61/TRAP translocons also associate with the oligosaccharyl transferase complex (OST) (Pfeffer et al. 2017); (Braunger et al. 2018) and recent findings suggest that TRAP also plays a role in coordinating the initial N-glycosylation process, which occurs in coordination with ER membrane translocation. Patients with germline mutations in different TRAP subunits have been described with aberrant protein glycosylation phenotypes (Losfeld et al. 2014); (Ng et al. 2015, 2019); (Dittner-Moormann et al. 2021) and a similar misglycosylation phenotype was observed upon specific TRAP subunit depletion experiments in cultured mammalian cells (Phoomak et al. 2021).

Here we present a single particle Cryo-EM reconstruction and an atomic model of the Sec61/TRAP translocon. The model reveals the general architecture of the entire heterotetrameric TRAP complex and suggests multiple interaction sites, both among the TRAP subunits and with different Sec61 subunits and the ribosome, which are consistent with the observed stable association. Our molecular dynamics (MD) simulations demonstrate that TRAP deforms the ER membrane in the vicinity of Sec61, which may assist in translocating specific Sec61 client polypeptides. TRAP contacts the ribosome in two locations, which can act to stabilize the ribosome exit tunnel for favourable polypeptide insertion, or may play a role in exerting the force required to perturb the local lipid environment. TRAPα contains a domain with homology to a bacterial chaperone which is positioned immediately below the Sec61 lumenal exit site and may form a direct binding site for inserting polypeptides that would prevent their back-diffusion. Proximity of TRAPα to where the OST has been visualized and the disease-association of TRAP in congenital misglycosylation diseases suggests a possible interplay with the glycosylation machinery. An atomic model of TRAP now allows future mechanistic studies to interrogate the interplay of TRAP and OST during protein N-glycosylation.

## RESULTS

### Cryo-EM model of the Sec61/TRAP translocon

During our work to characterize the structure of Sec61 bound to a substrate-selective cotransin analog (Rehan et al. 2022), we observed an additional density close to the Sec61 hinge and the Sec61γ subunit (**Fig. 1A**). The shape of the density in our single particle reconstruction closely resembles that of a published translocon-associated protein (TRAP) complex observed in cryo-electron tomography studies from isolated ER microsomes (Pfeffer et al. 2017). Western blot analysis of our isolated Sec61/ribosome preparations with a specific TRAPα antibody confirmed the presence of TRAP (**Fig. S1.1**), and features in our initial single particle reconstruction allowed unambiguous identification of all TRAP transmembrane domains.

**Figure 1.**
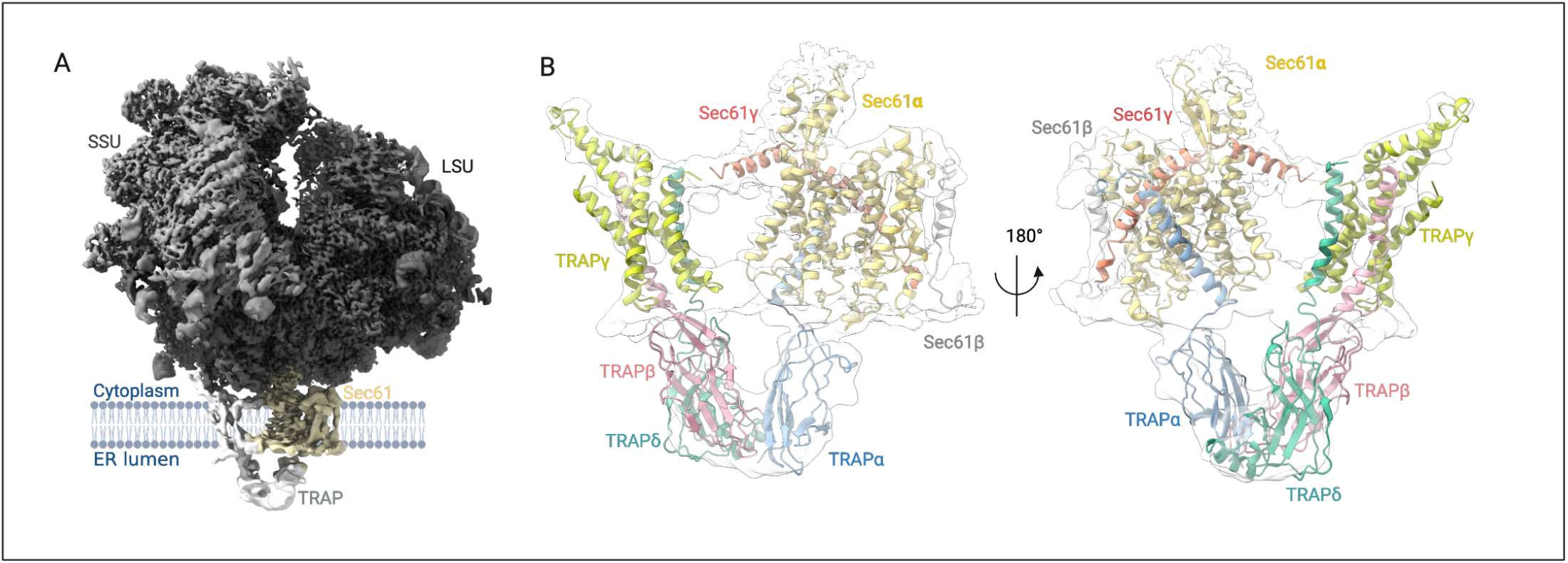
Cryo-EM structure of the Sec61/TRAP translocon. (A) Density of the mammalian 80S ribosome/Sec61/TRAP complex obtained after locally filtering the homogenous refinement output map in cryoSPARC (σ =0.6). Isolated complex subunits are shown for TRAP alpha (blue), beta (red), delta (green) and gamma (yellow). (B) Closeup of the Sec61/TRAP complex from the front (left) and back (right)

To refine the TRAP density, we used focused 3D classification to isolate TRAP-containing Sec61/ribosome particles, followed by signal subtraction to improve the local resolution of the TRAP subunits (**Fig. S1.2**). The extracted particles were further refined using heterogeneous and homogeneous 3D refinement tools in cryoSPARC to yield a reconstruction of the entire ribosome/Sec61/TRAP complex with an overall resolution of 2.6 Å. Resolution at the membrane region of Sec61 and TRAP varied between 4 and 6.5 Å, whereas resolution for the ER lumenal TRAP domains was limited to between 5.5 and 7.0 Å, presumably due to high mobility of this flexibly tethered unit (**Fig. S1.3**). The density for the TRAP complex was sufficiently detailed to build an atomic model of the TRAP subunits, which was not previously possible using the cryo-ET (Braunger et al. 2018). It should be noted that during data processing we only observe ribosome particles containing either Sec61 or Sec61/TRAP, but not Sec61 complexes with OST like observed before (Pfeffer et al. 2017); (Braunger et al. 2018), likely due to minor sequence differences between sheep and canine translocon components resulting in OST complexes being removed during our sample preparation.

As a starting point, we generated homology models for all four TRAP subunits using AlphaFold2 (Jumper et al. 2021) (**Fig. S1.4**). High confidence scores suggested that the entire TRAPγ subunit as well as the three TRAP lumenal domains would be valid models for assembling a model of the TRAP complex. Initial fitting of TRAPγ was unambiguous based upon clear densities for the TM and cytosolic helices which are in contact with ribosomal rRNA (**Fig. S1.5**). We also observed two additional TMs associated with the TRAPγ four-helix TM bundle, which we assigned as the single membrane anchors of TRAPβ and TRAPδ subunits.

We next used Alphafold2 Multimer (Evans et al. 2021) to model the TRAPα and TRAPβ complex. This provided a plausible and high confidence arrangement in which the two lumenal domains of TRAPα and TRAPβ form a roughly V-shaped arrangement. Placement of the TRAPα/β dimer into the observed density provided a good fit to the density and we conclude that this is likely reflective of the arrangement in the full TRAP complex. Additionally, we observed weak density at high contour levels for the disordered N-terminus of TRAPα, which further guided orientation of this domain in the map (**Fig. S1.6**). Initial placement of the TRAPδ lumenal domain was aided by a cryo-ET difference map comparing Sec61/TRAP complex isolated from normal or TRAPδ-deficient patient cells (Pfeffer et al. 2017) (**Fig. S1.7**). The resolution of the lumenal part of TRAP is limited, but a short alpha helix in the model of the TRAPδ lumenal domain suggested a plausible binding orientation relative to the TRAPα/β dimer. Promisingly, the dimer interface of the fitted TRAPα/β dimer agreed with an unbiased prediction by the HADDOCK 2.4 server (Roel-Touris et al. 2019). In the membrane part, we assign density for each TRAP transmembrane segment, including a long diagonal TRAPα TM that contacts the hinge loop of Sec61α and the backside of Sec61γ at the cytosolic side of the membrane. The C-terminal cytosolic region of TRAPα resides in proximity to ribosomal protein proteins uL26 and uL35, but is not visible in the cryo-EM map. After manual modeling, coordinatesfor Sec61, TRAP and protein subunits and RNA of the ribosomal large subunit were refined against the cryo-EM map in Phenix (Afonine et al. 2018).

To validate the Sec61/TRAP model, we used crosslinking mass spectrometry (XL-MS) (Kelly et al. 2022). Whereas the majority of the identified crosslinked peptide pairs are unsurprisingly observed between different ribosomal proteins, we do observe additional crosslinked peptide pairs between the TRAPβ and TRAPγ subunits, as well as between those and components of the 60S ribosome.

The most commonly identified crosslink, as well as two additional peptide pairs fit well within the expected crosslinking distance for DSS (10–30 Å) (data not shown).

### Architecture of the Sec61/TRAP translocon

In the cryo-EM structure, the macrocyclic cotransin inhibitor is bound to the Sec61 complex with an open lateral gate and a closed plug helix (Rehan et al. 2022). This is similar as observed in the cryo-ET study of the Sec61/TRAP complex in the ER membrane, where surprisingly the lateral gate was observed to be open in majority of Sec61 molecules (Pfeffer et al. 2017). The tetrameric TRAP complex binds to the backside of Sec61 on the opposite side of the lateral gate where nascent polypeptides insert into the lipid bilayer (**Fig. 2A**).

**Figure 2.**
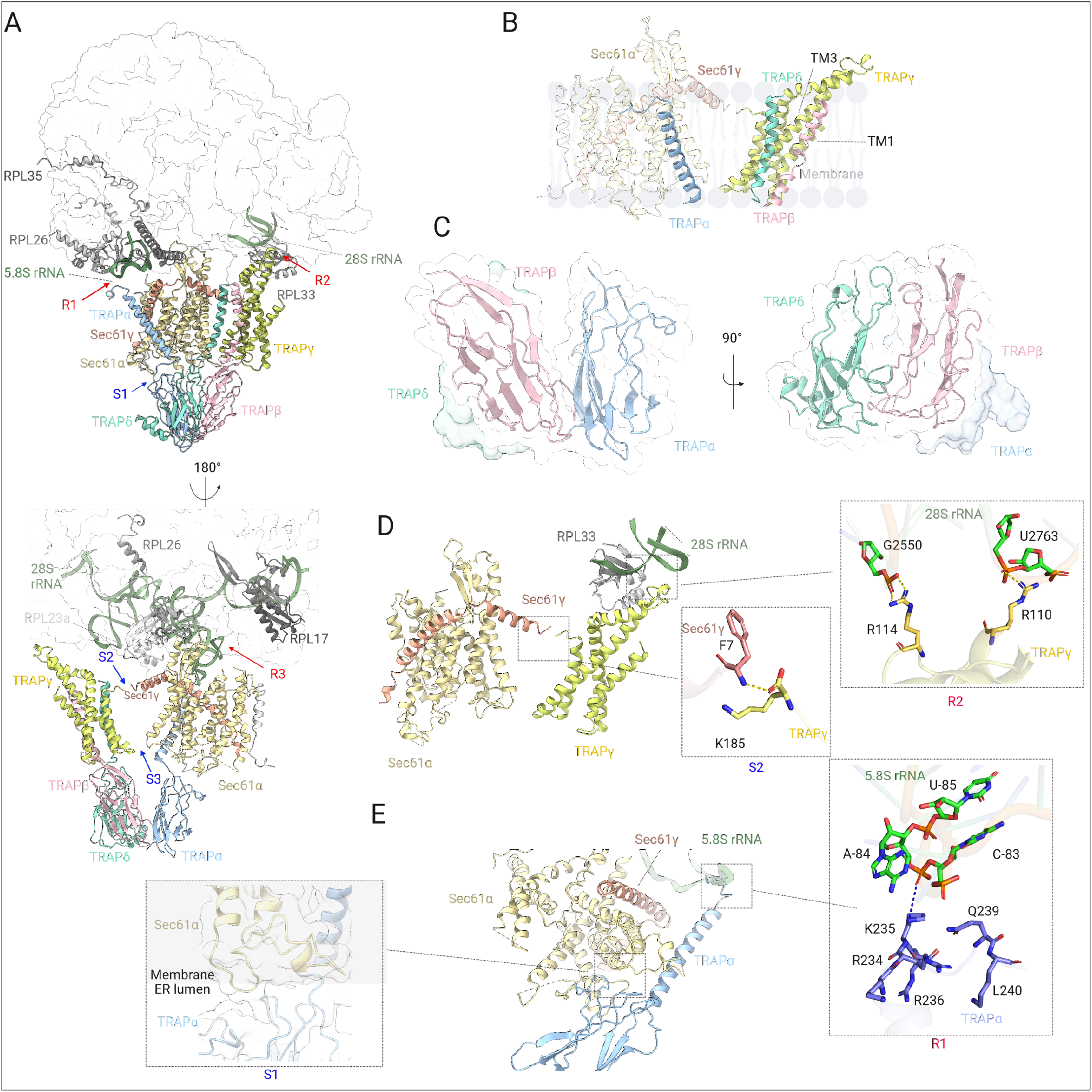
CryoEM structure of the TRAP complex and its interaction with Sec61 and Ribosome (A) Overview of the TRAP-Sec61-Ribosome complex structure highlighting the interaction sites of TRAP with Ribosome (R1, R2 and R3, indicated with red arrow), and TRAP with Sec61 complex (S1, S2 and S3, indicated with blue arrow), (B) Transmembrane domains of TRAPβ and TRAPδ interact with the transmembrane helices (TM1 and TM3) of TRAPγ to form a trimeric complex. TRAPα traverses the membrane diagonally away from the TRAPβ, TRAPδ and TRAPγ complex and forms a connection with the backside of Sec61γ on the cytosolic part of Sec61, (C) Formation of the trimeric TRAP complex between TRAPα, TRAPβ and TRAPδ in the lumenal region of the ER, (D) Interactions of TRAPα with Sec61α in the lumenal region (S1) and with the 5.8S ribosomal RNA in the cytoplasmic region (R1), TRAPα color-coded according to atom (nitrogen: blue, carbon: purple, oxygen: red), and 5.8S ribosomal RNA atoms are color-coded as, carbon: green, oxygen: red and nitrogen: blue). Hydrogen bond highlighted with the blue dashed line, (E) Interactions of Sec61γ with TRAPγ in the membrane region (S2), and TRAPγ with the 28S ribosomal RNA in the cytoplasmic region (R2),TRAPγ is color-coded according to atom as nitrogen: blue, carbon: yellow, oxygen: red), and 28S ribosomal RNA atoms are color-coded as, carbon: green, oxygen: red and nitrogen: blue), Sec61γ atoms as nitrogen: blue, carbon: light red, oxygen: red). Hydrogen bond highlighted with yellow dashed line. In all the figures (A-E), TRAP subunits colored as TRAPα:cyan, TRAPβ:pink, TRAPγ:green and TRAPδ: yellow, Sec61 complex colored as Sec61α:lightorange, Sec61β:grey and Sec61γ:red. 28S rRNA in light green, and 5.8S rRNA highlighted in green. All the ribosomal proteins highlighted in different shades of grey color.

TRAPγ forms a four-helix transmembrane bundle positioned in the backside of Sec61 in the membrane plane (**Fig. 2B)**, and the TRAPγ subunit resides predominantly in the membrane with helical section extending to the cytosolic side and contacting the ribosome (**Figs. 2A and 2D**). The N-terminal 30 residues of TRAPγ are predicted to form an alpha helix, but are not visible in our density and this region is presumably mobile or unstructured.

The transmembrane anchors of TRAPβ and δ form a bundle with TRAPγ TM helices, TM1 (Ser35–Arg49) and TM3 (Glu116–Ile156)(**Fig. 2B**). TRAPα, TRAPβ and TRAPδ subunits each contain a small folded beta sheet-rich lumenal domain that are each connected to the transmembembrane segments via flexible linker sequences (**Figs. 2A and 2C**). These three domains form a tight complex of ∼50×60×45 Å in the ER lumen and may contribute to the structural integrity of the TRAP complex. Importantly, the central TRAPβ subunit, sandwiched between TRAPα and TRAPδ lumenal domains has been shown to be critical for stability of the tetrameric TRAP complex (Phoomak et al. 2021) (**Fig.2C**). The TRAPα lumenal domain is positioned immediately below the central channel of Sec61α where nascent polypeptides emerge after Sec61 plug opening (**Figs. 2A and 2E**). TRAPα and TRAPβ are shown to be N-glycosylated in mammalian cells (Phoomak et al. 2021); (Wiedmann et al. 1987) and the glycosylation sites in our model (Asn136 and Asn191 in TRAPα and Asn107 and Asn91 in TRAPβ) are pointing away from the protein interaction sites (**Fig. S2.1**). TRAPα N-terminus is not visible in the density and is presumbaly unstructured; the approximately 80 N-terminal residues contain a putative Ca2+ binding motif, which is supported by our MD simulations (data not shown).

To resolve the key interactions among the various subunits of the Sec61/TRAP complex with the ribosome, we carried out atomistic molecular dynamics simulations of the entire Sec61/TRAP complex embedded in a lipid bilayer. We solvated the structure, and used two complementary force fields with the backbones of the RNA and proteins restrained, which revealed potential hydrogen bonding partners between the TRAP subunits (**Table S5**). At the TRAPα/β interface in the lumen, the Arg144–Glu114, was the only one with significant occupancy. In our model, these subunits have a relatively small interface (**Fig. 2C**), yet Ser76 of TRAPα and Glu17 of TRAPβ significantly contribute to the interface stability through an electrostatic interaction (**Table S6**). The lumenal interface between TRAPβ and TRAPδ is larger (**Fig. 2C**) with putative hydrogen bonds between Ile45–Asp82, Asn44–Asp82, and Ser27–Glu46. Additionally, the interface is stabilized by significant hydrophobic and electrostatic interactions by Pro84 and Asn30 of TRAPα and Arg79 of TRAPβ. Taken together, MD predicts the lumenal TRAPβ/δ interface to be significantly more stable than the lumenal TRAPα/β interface (**Table S6**), yet resolving the role of the highly charged and unstructured domain of TRAPα is beyond the sampling ability of present-day simulations. Apart from the lumenal domains, the N-terminal Ala173 of TRAPδ forms a hydrogen bond with Lys91 of TRAPγ at the cytosolic membrane interface. TRAPβ docks to TRAPγ both at the lumenal membrane interface (Arg49–Glu137) as well as within the membrane core (Ser159–Asn142). The Arg31– Glu150 hydrogen bond between TRAPβ and δ resides at the lumenal membrane interface.

The TRAPα lumenal domain is connected to a long 43-residue transmembrane helix, which interacts with the Sec61 lumenal hinge loop (**Fig. 2B and E**) and traverses the membrane diagonally forming a connection with backside of Sec61γ on the cytosolic domain of Sec61. At the lumenal interface, our MD analysis reveals Glu156 and Glu192 of TRAPα as potential hydrogen bonding partners with Arg205 and Tyr235 of Sec61α, respectively (Table S4). The Phe7 and Val8 residues of Sec61γ dock to Lys185 of TRAPγ at the cytosolic membrane interface (**Fig. 2D**), “**S2**”, whereas the Lys158 of TRAPγ forms a hydrogen bond with Asp357 of Sec61γ at the lumenal membrane interface (**Fig. 2D**), “**S3**”.

Sec61/TRAP binding to the ribosome is mediated by three interaction sites. First, the L6/L7 and L8/9 loops of Sec61α interact with with the ribosomal protein uL23 (Voorhees et al. 2014). In our model, The Sec61α residue Tyr416 forms a hydrogen bond with Ile156 of uL23, whereas both Ser408 and Gly403 of Sec61α can hydrogen bond to Glu84 of uL23 (**Table S7**) Sec61γ also participates in hydrogen bonding at this site, as Lys16, Arg20, and Arg24 of Sec61γ face Asp148 and the C-terminal Ile156 of uL23. The 28S ribosomal RNA residues C2526, G2433, and U2432 also form hydrogen bonds with Arg405, Arg273, and Lys268 of Sec61α, respectively. Second, the TRAPγ subunit also directly contacts the 28S ribosomal RNA through interactions involving TRAPγ Arg110 and Arg114 with 28S rRNA G2763 and G2550, respectively (**Fig. 2D**). The nearby Glu116 of TRAPγ hydrogen bonds to the ribosomal protein L38. Third, the cytosolic C-terminus of TRAPα, which is not visible in our density, contains residues that are positioned to interact with the 5.8S rRNA (**Fig. 2E**) and our MD simulations indicate an interaction of TRAPα Lys235 with the A84 of 5.8S rRNA (**Fig. 2E**).

A notable feature of the Sec61/TRAP complex is the large wedge-shaped cavity between Sec61 and TRAPβ/γ/δ with an approximate distance between the two complexes ranging between 11 and 39 Å (**Fig 2B**). In the ER membrane, this cavity is filled with lipids and it is possible that the local membrane environment between TRAP and Sec61 may affect the dynamics of Sec61 and thereby impact the kinetics of protein ER import and/or their cotranslational modification.

Overall, our structure reveals multiple interactions between the different TRAP subunits and also with subunits of the Sec61 complex, consistent with the stable biochemical nature of the TRAP complex and its tight binding to Sec61 (Hartmann et al. 1993); (Ménétret et al. 2008). The three interaction sites of TRAP and Sec61 with the ribosome likely provide a more stable complex that may may be required for translocation of TRAP dependent polypeptides.

We next probed the stability of this complex by unrestrained MD simulations of the Sec61/TRAP complex with interacting ribosomal proteins and RNA embedded in a membrane mimicking the composition of the mammalian ER membrane (**Fig. 3A**). Our atomistic microsecond-scale MD simulations based on the CHARMM36m/CHARMM36 force fields suggest that the anchoring interactions between Sec61, TRAP, and ribosome described above have a substantial effect on the stability of the Sec61/TRAP complex. The root mean squared deviation of the protein backbones demonstrates that the interaction with Sec61 alone does not stabilize TRAP, as TRAP displays a similar RMSD value of ∼20 Å whether together with Sec61 or alone. However, the anchoring of TRAPα and γ to the ribosomal proteins and RNA lead to a significant stabilization of TRAP and to an RMSD value of ∼10 Å (**Fig. 3B**). This effect was also verified with a complementary simulation using the Amber FF19SB/Lipid21/OL3 force fields (**Fig. S3E**). Unsurprisingly, the stabilization effect is the most substantial for TRAPα, which in the absence of the ribosome only anchors itself with Sec61α at the lumenal membrane interface, whereas the cytosolic end of the TM helix drifts towards the trimeric bundle of other TRAP subunits, leading to the loss of the characteristic V-shaped conformation. Still, due to the interactions between the lumenal domains, and the transmembrane bundle of TRAPβ, γ, and δ, the TRAP subunits do not dissociate during any of the 2 μs-long simulations.

**Figure 3.**
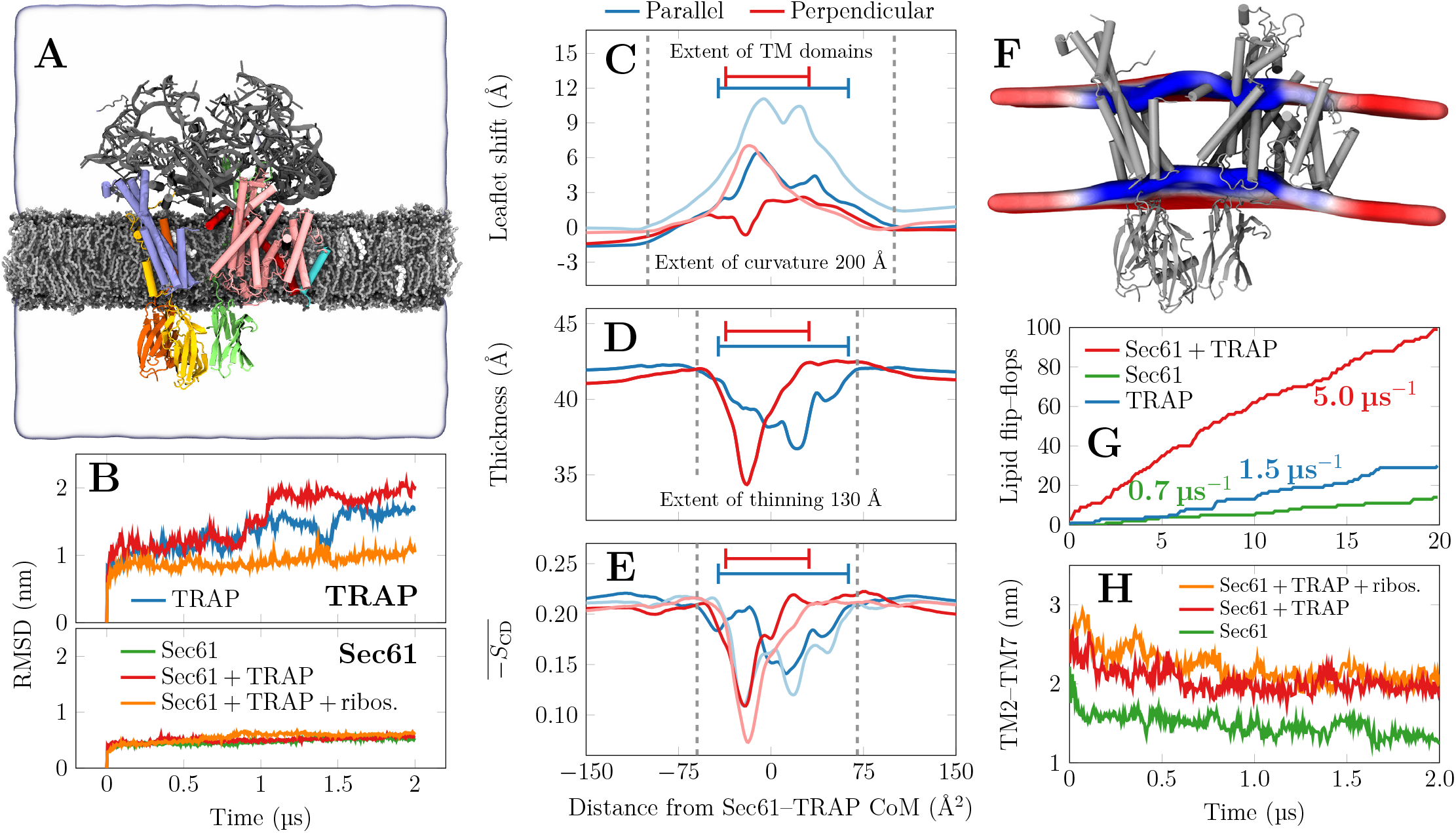
Membrane remodeling by the Sec61–TRAP complex as revealed by MD simulations. A) Snapshot of the initial conformation of the simulation system containing the Sec61/TRAP complex together with parts of the ribosome that interact with Sec61 or TRAP subunits. The distal parts of ribosome are restrained to model its large size without the need to model it entirely. TRAP subunits are shown in green (α), yellow (β), blue (γ), and orange (δ), whereas Sec61 subunits are shown in pink (α), cyan (β), or red (γ). The ribosomal proteins and RNA fragments included are drawn in gray, The lipids are shown in silver with gray head groups, and cholesterol in white. The extent of the simulation cell is highlighted by the transparent surface. The lipid hydrogens, water molecules, and ions are not rendered for clarity. B) Root mean square deviation (RMSD) of the TRAP and Sec61 structures when simulated in different assemblies. Sec61 is always stable, yet TRAP conformation shows significant variations in the absence of ribosomal anchoring. C) Quantitative characterization of membrane perturbations using g_lomepro Gapsys et al. (2013). The vertical shift of the lipid phoshoprus atoms. Profiles are calculated parallel to the axis connecting Sec61 and TRAP and perpendicular to it. Darker lines show the upper (cytosolic) leaflet and lighter ones the lower (lumenal) leaflet. The extent of the protein TM regions is highlighted. D) Membrane thickness calculated as the difference between the phoshorus profiles of the two leaflets in C). E) Local membrane ordering calculated as the average of the deuterium order parameters of carbons 2–15 in the palmitate chains of phosholipids. F) The perturbation of the membrane in the simulation containing Sec61/TRAP anchored by the ribosome contacts. The average positions of the phosphorus atoms is shown by the coloured surface cut at the protein location. The colour depicts local thickness, ranging from 37 Å (blue) to 43 Å (red) (average of the protein-free control simulation is 31.8 Å). G) Lipid flip–flops as a proxy to membrane perturbation and permeabilization. The cumulative POPC flip–flops in the coarse-grained simulations. In simulations with individual TRAP subunits, no flip–flops were observed, but they are promoted by the bundle of TRAPβ, TRAPγ, and TRAPδ TM domains. Sec61 alone has a minor effect, but together with TRAP the lensing effect significantly accelerates flip–flops. H) The distance of the lateral gate helices TM2 and TM7 in the atomistic simulations. The presence of TRAP seems to help maintain the gate in a more open conformation.

### TRAP binding alters the local membrane environment and stabilizes Sec61 in a conformation with an open lateral gate

To interrogate the effects of TRAP association for the ER lipid membrane and provide insights for consequent effects for the Sec61 translocon, we analyzed our atomistic MD simulations based on the CHARMM36/CHARMM36m force fields. During the 2 μs simulations containing the Sec61/TRAP complex and proximal ribosome components (**Fig. 3A**), we observed a dramatic perturbation of the ER membrane in the vicinity of the Sec61/TRAP transmembrane regions (**Fig. 3B**). In particular, the local membrane structure budges towards the cytosol (**Fig. 3C**), and the curved region ranges over a distance of ∼200 Å, spanning an effective area extending well beyond the Sec61/TRAP complex. We note that the membrane deformation effect is more pronounced in parallel to the axis connecting Sec61 and TRAP than perpendicular to it (**Fig. 3C**). Moreover, the ER lumenal membrane leaflet is perturbed to a larger extent with a local bulge reaching ∼10 Å, which leads to membrane lensing, i.e. the simultaneous curving and thinning of the membrane (**Fig. 3D**). This lensing effect is brought about by membrane remodelling to accommodate the TRAP and Sec61 TM domains while tilted to adopt a V-shaped formation through interactions with the ribosome. We verified the lensing effect with complementary atomistic and coarse-grained simulations based on the Amber FF19SB/Lipid21/OL3 and Martini 3 force fields, respectively, and the lensing effect was systematically reproduced (**Fig. S3C and D**).

The distribution of local membrane thicknesses can be fitted by two Gaussians corresponding to values of 41.7±0.58 Å and 38.1±2.37 Å. The former agrees extremely well with the sharp single Gaussian distribution of our protein-free control system (41.8±0.57 Å) and thus corresponds to an unperturbed region. The smaller value corresponds to thinner regions and comes with a broad distribution, highlighting the diversity in local lipid environments around the Sec61/TRAP complex. The local thinning and curving may sort ER lipids in a specific manner around the Sec61/ TRAP complex, however our attempts to evaluate the effects of curvature on local ER lipid composition using MD simulations did not display different partitioning preferences in a curved ER membrane. We note that our simulation system consisted of identical acyl chain lengths and more detailed lipidomics studies may be required to better understand the curvature–composition coupling that might play a role on the dynamics of the translocon.

The membrane lensing in the Sec61/TRAP complex leads to differences in lipid packing across the leaflets, and the profiles of mean deuterium order parameters of the palmitate chains reveal that the ER lumenal leaflet is more disordered. The differences become pronounced in the vicinity of the bundle formed by TRAPβ, TRAPγ, and TRAPδ, where the lipids in the cytosolic leaflet show modest perturbation, while those in the lumenal leaflet are significantly disordered (**Fig. 3E**). The disordering effect spans the dimensions of the entire Sec61/TRAP complex and the order parameter distributions in both leaflets can be fitted by two Gaussians. Both leaflets display a narrow distribution around S_CD_ ≈0.20, in agreement with the protein-free control system. The more disordered component in the cytosolic leaflet has S_CD_=0.15±0.04, whereas the lumenal leaflet comes with a broader distribution with S_CD_CD=0.19±0.09.

The periodicity of the simulation cell restricts the development of curvature in typical membrane protein simulations. We therefore also embedded the Sec61/TRAP complex with the ribosome in a lipid bicelle with a diameter of ∼20 ns which can curve to any degree (Kluge et al. 2022) (**Fig. S3A**). In this unrestrained system, the bicelle demonstrated a more pronounced lensing effect and a curvature of ∼285 Å− 1 (**Fig. S3B**).

Next we set out to study what are the minimum components required to produce the lensing effect by performing control simulations of TRAP alone, Sec61 alone, and the Sec61/ TRAP complex in the absence of the ribosome. Sec61 and TRAP were unable to induce any curvature alone. Curiously, we did not detect any significant membrane curvature even with the Sec61/TRAP complex in the absence of ribosomal anchoring, although thinning of this flat membrane was observed to a similar degree as in the curved one with ribosomal anchoring (**Fig. S3H**,**I**). These findings suggest that the observed lensing requires the presence of the V-shaped Sec61/ TRAP complex and its anchoring to the ribosome.

The TM domain of TRAPβ contains two prolines (P158 and P163) that break the α helix, and MD simulations suggest that these prolines, together with the nearby N35, N141, and N142 residues of TRAPγ, attract water into the membrane and facilitate permeation events through the ER membrane. We used coarse-grained simulations with the recent Martini 3 force field (Souza et al. 2021) to investigate at longer time scales. We chose to model ribosomal anchoring by restraining the Sec61 and TRAP backbones. During a 20 μs simulation of this complex embedded in a POPC bilayer, we observed ∼100 spontaneous lipid flip–flop events to take place in the vicinity of the transmembrane trimeric bundle formed by TRAPβ, TRAPγ, and TRAPδ. (**Fig. 3G**). We then performed simulations of all the TRAP subunits alone, yet observed no flip–flops, highlighting the role of the trimeric bundle. Curiously, simulations with all TRAP subunits present demonstrated a significantly smaller number of flip–flops than when TRAP was paired with Sec61, yet this difference could not be explained by the flip–flops promoted by Sec61. Thus, we hypothesize that the permeability is enhanced by the local membrane thinning, curving, and disordering that lead to its increased fluidity.

Finally, we sought to understand what role TRAP and ribosome binding and the associated membrane perturbation may have for Sec61 conformational dynamics. We assessed the effect of TRAP and ribosome for lateral gate dynamics by measuring changes in distance across Sec61 lateral gate helices TM2 and TM7 over time (**Fig. 3H**). The simulations were initiated starting from an open conformation observed in our structure of Sec61 primed with a macrocyclic cotransin inhibitor (Rehan et al. 2022). In simulations without TRAP, we observed the lateral gate closing rapidly, whereas simulations with TRAP included retained an open lateral gate conformation. Finally, anchoring the of the Sec61/TRAP complex to the ribosome resulted in the lateral gate opening even further. To verify the robustness of the lateral gate observations, we analyzed our control simulation of Sec61/TRAP complex together with the ribosomal subunits using atomistic Amber force fields. In this simulation, the lateral gate initially closed during the equilibration stage of the simulations, yet reopened after ∼1.7 μs of the production simulation, which supports our notion that TRAP and ribosome association promotes the open gate of Sec61, which was also observed in an earlier cryo-ET study of Sec61/TRAP in intact ER (**Fig. 3F,G**).

## DISCUSSION

The evolutionarily conserved heterotrimeric Sec61 translocon is sufficient for ER translocation of select secretory polypeptides across the ER membrane. However, in mammalian cells many, if not most, proteins require additional auxiliary components to efficiently translocate certain secretory polypeptides (Fons et al. 2003); (Hegde et al. 1998); (Conti et al. 2015). Cryo-electron tomography has shown that the predominant stable form in which Sec61 exists in ER membranes is in complex with the heterotetrameric TRAP complex (Pfeffer et al. 2017). Biochemically, the TRAP complex forms a stable constitutive complex with Sec61 and promotes ER insertion of specific secreted and integral membrane proteins by a yet unidentified mechanism (Fons et al. 2003); (Hartmann et al. 1993). Here, we present a single particle cryo-EM structure of the TRAP complex bound to the mammalian Sec61 translocon and the ribosome. Our structure shows that TRAP binds to the ribosome through contacts to the 5.8S and 23S rRNAs at two sites in addition to the Sec61α L6/L7 loop (Voorhees et al. 2014). Our molecular dynamics simulations indicate that TRAP association enhances the perturbation of the local membrane environment surrounding Sec61, which we propose is required for lateral gate engagement of inefficient signal peptides and transmembrane segments. The V-shaped conformation of TRAP remodels the lumenal leaflet of the ER membrane and leads to significant local curvature and thinning, which can further modulate the structure and dynamics of the Sec61 channel. The TRAPα lumenal domain is situated immediately below the Sec61 channel and nascent polypeptide binding to this domain may assist by biasing diffusion into the ER lumen.

While TRAP dependency cannot be predicted from client protein sequence alone, a proteomics study has suggested that specific features of the nascent signal peptide are required and especially sequences with low hydrophobicity and that are enriched in proline and glycine residues would be over-represented among TRAP client proteins (Nguyen et al. 2018). Further, it has been shown that the signal peptides of TRAP clients such as prion protein or insulin are a key determinant for TRAP engagement at an early stage of protein insertion (Kriegler et al. 2020b,a); (Fons et al. 2003). Force pulling experiments suggest that the signal peptides of TRAP clients cannot efficiently intercalate between lateral gate helices when TRAP is depleted (Kriegler et al. 2020b). Our MD results now suggest that the presence of TRAP changes the conformational landscape of the Sec61 lateral gate, which is consistent with inability of specific inefficient signal peptides to engage with the lateral gate. The effect of TRAP could be mediated either by conformational stabilization through direct Sec61–TRAP interactions or via the membrane through a hydrophobic mismatch mechanism (Killian 1998); (Yeagle et al. 2007) coupled with local membrane curvature. Moreover, the local curvature and thinning could also sort the ER lipids, resulting in a specific local lipid composition. Notably, the stiffening of ER membranes by loading them with excess cholesterol has been shown to inhibit Sec61-mediated membrane translocation in biochemical experiments (Nilsson et al. 2001), which further supports the notion that membrane fluidity and deformability may play an important role in tuning Sec61 translocation activity.

After initial Sec61 engagement of human prion protein, a second TRAP-dependent force pulling event has been described (Kriegler et al. 2020b), which presumably occurs in the ER lumen and may represent direct binding to the TRAPα lumenal domain that is situated approximately 20 Å away from the lumenal end of the Sec61 channel. Further, it was shown that a hydrophilic sequence immediately following the prion protein signal peptide is required for the second force pulling event. TRAP has also been shown to be required for establishment of correct topology of an integral multipass model membrane protein with mildly hydrophobic N-terminal TM segments (Sommer et al. 2013). Because the unstructured N-terminal tail of TRAPα is highly negatively charged and positioned close to the Sec61 lumenal end, we speculate that this charged flexible segment may attract hydrophilic peptide sequences that initially emerge in the ER lumen but do not have a strong propensity to obtain the correct topology for example because of a lack of cytosolic positively charged residues. At an early stage of ER insertion before a significant length of nascent chain has inserted into the ER lumen, many nascent polypeptides can readily diffuse back into the cytosol resulting in loss of ER insertion and instead targeting the mislocated polypeptide for cytosolic degradation. Because the structure of the TRAPαlumenal domain resembles that of bacterial chaperones, it it possible that many TRAP client proteins directly interact with this domain at a site immediately below the Sec61 channel where inserting polypeptides emerge into the ER lumen. Such an interaction with hydrophilic sequences downstream of the signal peptide or transmembrane segment would bias polypeptide diffusion in the direction of the ER lumen before a downstream hydrophobic sequence can engage with lumenal chaperones. The positioning of this domain in our structure, the abundance of charged residues in the TRAPα folded domain and un structured tail segment Hartmann and (Prehn 1994) and the observed crosslinks between nascent polypeptides and TRAP subunits (Wiedmann et al. 1987) support the notion of direct TRAPα interactions with inserting polypeptides in the ER lumen.

In addition to ER translocation, most secretory proteins require correct processing for accurate folding and functionality, and TRAP has been implicated in influencing secretory protein modification. A recent study demonstrated that depletion of TRAP subunits strongly reduced insulin biogenesis and processing in human beta-cells, whereas re-expression of the missing subunits restored expression, signal peptide processing and disulphide bond formation (Huang et al. 2021); (Li et al. 2019). Preproinsulin is a short polypeptide which is assumed to be targeted to Sec61 at least partially in a post-translational manner, and the observed TRAP dependence may be explained by a requirement to engage with TRAPα in the ER lumen to prevent the polypeptide slipping back into the cytosol following signal peptide cleavage and termination of polypeptide synthesis. Further, several studies suggest that TRAP may play an important role in directing cotranslational N-glycosylation of nascent polypeptides, which is carried out by the STT3A oligosaccharyl transferase (OST) complex that is situated in the immediate proximity of TRAP α and δ subunits (**Fig. 4A**). Several studies have identified germline TRAP subunit mutations in patients with protein misglycosylation defects (Losfeld et al. 2014); (Dittner-Moormann et al. 2021), and another study identified TRAP as a new factor required for correct cell-surface N-glycosylation in mammalian cells (Phoomak et al. 2021). Although the effects of TRAP depletion on glycosylation can in principle be explained by prevention of protein entry into the ER lumen, a more direct role is also possible. The proximity of the TRAPα and δ subunits to OST and its catalytic active site (**Fig. 4B**), suggest that nascent polypeptide binding to TRAPα could prevent unwanted backdiffusion when only a short polypeptide has reached the ER lumen around the time when signal peptide cleavage occurs and transient binding of specific polypeptide sequences to TRAPα could position a glycosylation sequence motif in an optimal configuration for N-glycan addition to occur.

**Figure 4.**
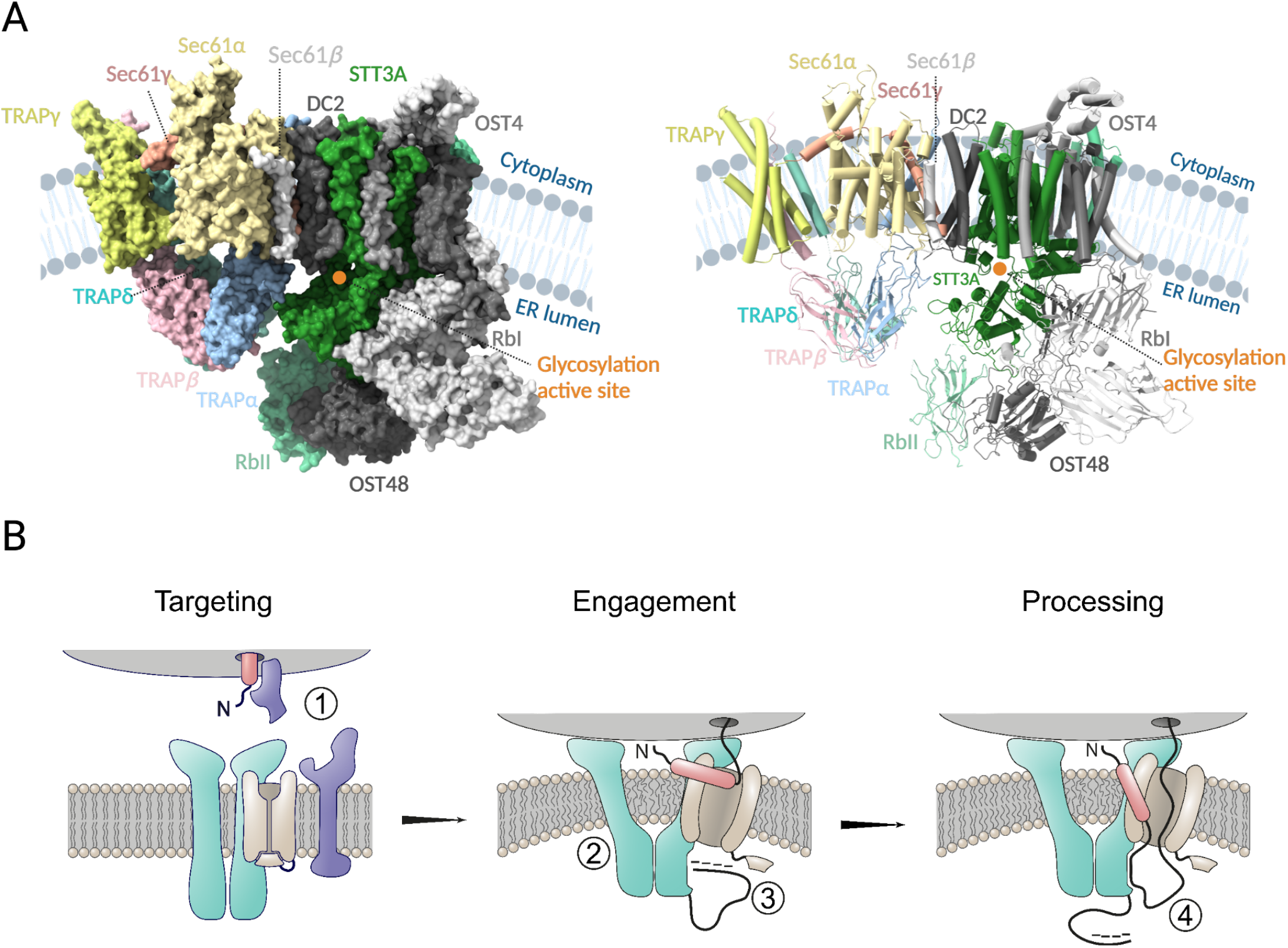
Functional model of the role of TRAP in nascent polypeptide processing. (A) Structure of the TRAP and OST-A (PDB ID: 6S7O) modeled in the cryoET density of Sec61/TRAP and OST complex (emd: 3068), surface (right) and cartoon (left) representation and .TRAP subunits colored as TRAPα:cyan, TRAPβ:pink, TRAPδ:green and TRAPγ: yellow, Sec61 complex colored as Sec61α:lightorange, Sec61β:grey and Sec61γ:red. Most of the OST subunits colored in grey except STT3A and RbII lie in proximity to the lumenal domains of the TRAP complex and are colored green and light green, respectively. The glycosylation active site of the STT3A domain is highlighted with an orange circle. (B) Targeting of the ribosome–nascent complex is carried out by SRP and SRP receptor (1). Docking of the ribosome induces a conformational change in the TRAP/Sec61 complex, resulting in membrane perturbation which increases the fluidity of the local lipid environment about the Sec61 lateral gate (2). After successful lateral gate engagement and plug displacement, nascent polypeptide is exposed to the ER lumen, where transient interactions with the negatively charged TRAPα flexible N-terminal loop may encourage correct topology and complete lateral gate intercalation of the signal peptide (3). The TRAPα lumenal domain is proximal to the lumenal exit of Sec61, and it may directly bind the nascent polypeptide, serving to prevent backdiffusion of problematic polypeptide sequences (4). Finally, interaction with TRAPα lumenal domain may also serve to transiently restrict the nascent polypeptide for presentation to downstream processing events.

The endoplasmic reticulum is a key site of protein biogenesis, and in highly proliferative or secretory cells the protein flux through the ER poses a significant challenge to the protein biogenesis machinery. In addition, the ER is a major site of lipid metabolism in cells and perturbations such as viral infection, change in redox environment or hypoxia can cause dramatic changes to the ER lipid composition with downstream effects to ER protein quality control. Local ER membrane perturbation, especially membrane thinning, has been suggested to lower the energetic barrier for transmembrane segment insertion and extraction (Wu et al. 2020). Our work now suggests that alterations to the ER membrane may also impact Sec61-mediated protein insertion and highlights the importance of understanding the localized membrane effects resulting from Sec61 cofactor association. Further, different natural and synthetic small molecule inhibitors of Sec61 are accommodated within Sec61 in subtly different open conformations (Rehan et al. 2022); (Itskanov et al. 2022) and detailed understanding of Sec61 conformational control by the surrounding lipid and protein environment will be important for targeting Sec61 with therapeutic small molecules. Further structural and mechanistic work is required to understand a possible direct role that TRAP may have in directing protein N-glycosylation with OST.

## EXPERIMENTAL PROCEDURES

### METHODS

#### Isolation of Sec61 complexes for Cryo-EM and data collection

RNC–Sec61–TRAP complexes were purified as described earlier (Rehan et al. 2022).). Briefly, 50 μl of sheep rough ER microsomes (SRM) were thawed and solubilized in 1% LMNG detergent with occasional mixing on ice for 1 h. Insoluble material was separated by centrifugation at 22000 rpm for 15 min. Clarified supernatant was loaded on a 1 ml Superose-12 gel filtration column pre-equilibrated with a buffer containing 0.003% LMNG. 10 fractions each containing 100 μl samples were collected and absorbance at A260 was measured using nanodrop spectrophotometer. Final concentration of the sample was estimated using the molar extension coefficient of eukaryotic ribosomes (Voorhees et al. 2014). Sample was centrifuged at 22000 rpm for 10 min to get rid of any aggregates before freezing grids. Purified sample was analyzed on western blot using anti-Sec61α, Anti-TRAPα and anti-RPL18 antibodies.

#### Data processing

Cryo-EM data processing was performed with Relion 3.046 (Zivanov et al. 2018) maintained within the Scipion 3.0.7 software package (Sharov et al. 2021), and also with cryoSPARC v3.3.2. A total of 1,089,031 particles were picked from 30,230 motion-corrected micrographs with SPHIRE-crYOLO47, contrast transfer function (CTF) parameters were estimated using CTFFIND448, and 2D and 3D classifications and refinements in Scipion were performed using RELION49. A total of 2,66,968 selected particles contributing to the best 3D classes were subjected to iterative rounds of 3D refinement until the FSC converged at 3.5 Å. The output particles from refinement were then 3D 9 of 17 classified without alignment, which generated ten classes with clearly distinguishable translating and non-translating ribosomes. To preclude density contributions from nascent polypeptides, only non-translating ribosome–Sec61 complexes were submitted to iterative CTF refinement, resulting in a final map that resolved to 3.2 Å resolution. To refine the TRAP density, 3D focused classification was performed to identify TRAP-containing particles, followed by signal subtraction the best class with 93,857 particles containing clear density of Sec61/TRAP complex (Fig. S2.2). In cryoSPARC, ab initio reconstitution generated two volumes with a clear density of Ribosome, TRAP and Sec61 (Fig. S2.2). Further heterogeneous 3D refinement generated volumes with FSC (0.143) converged at 3.0 Å resolution (61177 particles) and 3.2 Å resolution (29142 particles). Homogeneous 3D refinement was performed using the obtained volumes from heterogeneous 3D refinement that generated high-resolution 3D maps with FSC (0.143) converged at 2.7 Å (61177 particles) and 2.9 Å (29142 particles) resolution. Obtained map with 2.7 Å resolution showed a better density of the Sec61/TRAP complex in the lumenal and membrane region.

#### Model building and refinement

Initial models of the sheep TRAP subunits were built using AlphaFold2 (AF2) (Jumper et al. 2021) developed by DeepMind that predicts a protein’s 3D structure from amino acid sequence. Amino acids sequences of TRAP subunits with UniProt sequence IDs, A0A6P7D666 for TRAPα, A0A6P3TVC6 for TRAPβ, W5NYA9 for TRAPγ and W5P940 for TRAPδ were used for structure modeling. The dimer model of the TRAPα and TRAPβ was generated using ColabFold (Mirdita et al. 2022). The obtained AF2 models were initially fitted into the cryoEM density using ChimeraX (Goddard et al. 2018). The model was further refined in Phenix (Afonine et al. 2018). Further, the final model was obtained by several rounds of rebuilding in COOT (Casañal et al. 2020) and refinement in Phenix (Afonine et al. 2018).

#### Molecular Dynamics Simulations

We performed a vast set of molecular dynamics (MD) simulation to study the structure, dynamics, and interactions of the Sec61–TRAP–ribosome complex. These were based either on atomistic CHARMM (CHARMM36 & CHARMM36m)(Klauda et al. 2010; Huang et al. 2017; Denning et al. 2011) or Amber (FF19SB, Lipid21, OL3) and (Tian et al. 2019; Dickson et al. 2022; Zgarbová et al. 2011)) force fields, or the coarse-grained Martini 3 (Souza et al. 2021) force field. First, using atomistic MD simulations of the complex embedded in a lipid bilayer and with the protein backbone restrained, we refined the side chain conformations and extracted information on hydrogen bonding partners and other key interactions between the different protein subunits. Secondly, using atomistic MD simulations of the ER membrane-embedded Sec61– TRAP–ribosome complex, the Sec61–TRAP complex, Sec61 alone, or TRAP alone, we studied the effects of the inter-subunit interactions on the complex structure and stability. Additionally, we resolved the effects of the protein assemblies on the structure of the host membrane, namely its thickness, acyl chain order, and curvature. A protein-free ER membrane was used as a control. Further evidence for membrane remodeling was obtained by simulating the Sec61–TRAP–ribosome complex in a bicelle, as well as through coarse-grained simulations of the complex. Details on the setup of the simulation systems, their composition, the used simulation parameters, and the performed analyses are available in the SI.

#### Crosslinking of the RNC-Sec61-TRAP complexes

Three different concentrations (1 mg/mL, 1.2 mg/mL and 2 mg/mL) of the RNC–Sec61–TRAP complexes in 50 mM HEPES, pH 7.4, 300 mM KOAc, 10 mM MgOAc, 1 mM DTT and 0.003% LMNG were crosslinked with 0.5 and 1.0 mM heavy/light DSS (DSS-H12/D12, Creative Molecules Inc., 001S), respectively. Non-crosslinked samples were kept as controls. All samples were incubated for 60 min at 25°C, 1000 rpm. The cross-linking reaction was quenched with a final concentration of 50 mM of ammonium bicarbonate for 15 min at 25°C, 1000 rpm.

#### Sample preparation for mass spectrometry

All samples were precipitated by trichloroacetic acid, and the precipitated proteins washed with acetone. The precipitated proteins were denatured using an 8 M urea– 100 mM ammonium bi-carbonate solution. The cysteine bonds were reduced with a final concentration of 5 mM Tris (2-carboxyethyl) phosphine hydrochloride (TCEP, Sigma, 646547) for 60 min at 37°C, 800 rpm and subsequently alkylated using a final concentration 10 mM 2-iodoacetamide for 30 min at 22°C in the dark. For digestion, 1 μG of lysyl endopeptidase (LysC, Wako Chemicals, 125-05061) was added, and the samples incubated for 2 h at 37°C, 800 rpm. The samples were diluted with 100 mM ammonium bicarbonate to a final urea concentration of 1.5 M, and 1 μG of sequencing grade trypsin (Promega, V5111) was added for 18 h at 37°C, 800 rpm. The digested samples were acidified with 10% formic acid to a final pH of 3.0. Peptides were purified and desalted using C18 reverse phase columns (The Nest Group, Inc.) following the manufacturer’s recommendations. Dried peptides were reconstituted in 10 μL of 2% acetonitrile and 0.1% formic acid prior to MS analysis.

#### Liquid chromatography tandem mass spectrometry

A total of 2 μL of peptides were analyzed on an Orbitrap Eclipse mass spectrometer connected to an ultra-high-performance liquid chromatography Dionex Ultra300 system (both Thermo Scientific). The peptides were loaded and concentrated on an Acclaim PepMap 100 C18 precolumn (75 μm × 2 cm,) and then separated on an Acclaim PepMap RSLC column (75 μm × 25 cm, nanoViper,C18, 2 μm, 100 Å) (both columns Thermo Scientific), at a column temperature of 45 °C and a maximum pressure of 900 bar. A linear gradient of 2% to 25% of 80% acetonitrile in aqueous 0.1% formic acid was run for 100 min followed by a linear gradient of 25% to 40% of 80% acetonitrile in aqueous 0.1% formic acid for 20 min. One full MS scan (resolution 120,000; mass range of 400-1600 *m*/*z*) was followed by MS/MS scans (resolution 15,000) of the 20 most abundant ion signals. Precursors with a charge state of 3-8 were included. The precursor ions were isolated with 1.6 *m*/*z* isolation window and fragmented using higher-energy collisional-induced dissociation (HCD) at a normalized collision energy (NCE) of 30 (all samples), or stepped NCE of 21, 26, 31 (RNC-Sec61-TRAP complexes at 2 mg/mL). The dynamic exclusion was set to 45 s.

#### Cross-linking data analysis

All spectra from cross-linked samples were analyzed using pLink 2 (version 2.3.10). To keep the search space limited, the target protein database contained the sequence for the Ovis aries Sec61 and TRAP complexes, as well as those of the 60S ribosome only. pLink2 was run using default settings for conventional HCD DSS-H12/D12 cross-linking, with trypsin as the protease and up to 3 missed cleavages allowed. Peptides with a mass range of 600–6000 *m*/*z* were selected (peptide length 6–60 residues) and the precursor and fragment tolerance were set to 20 and 20 ppm, respectively. The results were filtered with a filter tolerance of 10 ppm and a 5% FDR.

## ACKNOWLEDGEMENTS

Research was supported by funding: VOP from the Academy of Finland (338836 and 314672), the Sigrid Juselius Foundation and the Jane and Aatos Erkko Foundation. JTH from the Academy of Finland (314669). M Javanainen from the Academy of Finland (338160). We are thankful to Pasi Laurinmaki and Benita Löflund (CryoEM unit, Institute of Biotechnology, University of Helsinki) for the technical support. We thank Jason van Rooyen (Beamline scientist, Diamond Light Source, UK) for the data collection. Support from the Swedish National Infrastructure for Biological Mass Spectrometry (BioMS) and the SciLifeLab, Integrated Structural Biology (ISB) platform is gratefully acknowledged (L.H.). We thank CSC–IT Center for Science for computational resources.

