## Supplemental Information File for "Molecular view of ER membrane remodeling by the Sec61/TRAP translocon"

### Supplementary Material for: Molecular view of ER membrane remodeling by the Sec61/TRAP translocon

September 30, 2022

### Contents


S1 Supplementary Methods . . . . . S2
S1.1 Molecular Dynamics Simulations . . . . . S2
S2 Supplementary Results . . . . . S8
S2.1 Purification of the Sec61/TRAP translocon . . . . . S8
S2.2 CryoEM data processing workflow . . . . . S10
S2.3 Local resolution . . . . . S11
S2.4 AlphaFold models . . . . . S12
S2.5 Fit of the individual TRAP subunit models in the cryo-EM density . . S13
S2.6 Density of the N-terminal TRAP$\alpha$ . . . . . S14
S2.7 Fit of the Sec61/TRAP atomic model to the cryo-ET density . . . . . S15
S2.8 Glycosylation sites on TRAP$\alpha$ and TRAP$\beta$ . . . . . S16
S2.9 Refinement and model statistics . . . . . S17
S2.10 Supplementary results from MD simulations . . . . . S18
S2.11 Hydrogen bonding between Sec61 and TRAP . . . . . S21
S2.12 Hydrogen bonding between TRAP subunits . . . . . S22
S2.13 Key interactions between TRAP subunits . . . . . S23
S2.14 Hydrogen bonding between Sec61/TRAP and the ribosome . . . . . S24

### S1 Supplementary Methods

#### S1.1 Molecular Dynamics Simulations

We performed an extensive set of molecular dynamics simulations for the Sec61–TRAP–ribosome complex and its various subcomplexes using both atomistic and coarse-grained simulations. These simulations were

1. Atomistic simulations of the Sec61–TRAP–ribosome complex in a lipid bilayer with backbone restraints on the protein and RNA backbones to resolve the key hydrogen bonding partners, and the interactions stabilizing the lumenal domain arrangement of TRAP.

2. Atomistic simulations of fully dynamic Sec61–TRAP–ribosome in a lipid bilayer to resolve the conformational stability of this complex and its various subcomplexes, and to analyze the effect of these complexes on the membrane structure.

3. Atomistic simulations of the Sec61–TRAP–ribosome complex in a bicelle to analyze its effects on membrane curvature.

4. Coarse-grained simulations of the TRAP–Sec61 complex in a lipid bilayer with backbone restraints to study the effect of this complex on membrane structure and on lipid flip–flops.

Our model followed the deposited PDB; certain Sec61$\alpha$ loops could not be rigorously modeled, so they were truncated yet continuous in the simulation model. Moreover, the unstructured part of the lumenal domain of TRAP$\alpha$ were omitted.

###### All-atom simulations for interaction analysis

We embedded the final atomistic model consisting of Sec61, TRAP, and part of the ribosome (in the vicinity of Sec61 and TRAP subunits) into a POPC bilayer. The system was hydrated, and neutralizing $K^+$ ions were added together with 144 mM of KCl. The membrane contained a total of 2000 lipids and 200 waters per lipid (400,000 in total), spanned dimensions of $\sim 26 \times 26 \times 23$ nm$^3$, and contained 1.58 million atoms. The model was set up using the CHARMM-GUI web portal [18, 34]. Two complementary force fields were used: Of the CHARMM family, the protein was described by the CHARMM36m force field [16], the lipids with CHARMM36 [20], and RNA with CHARMM36 [8]. Water was described with the CHARMM-specific TIP3P model [19, 10]. Of the Amber family, we chose the FF19SB force field [32] for the protein, the Lipid21 force field [9] for the lipids, and the standard TIP3P model [19] for water.

The structures and force field files were downloaded in GROMACS-compatible formats [21]. The standard CHARMM-GUI equilibration protocol was performed for both systems with the restraints on protein backbone and sidechains increased. Moreover, the backbone restraints were maintained throughout the equilibration steps. Finally, the protein backbone atoms were restrained by a force constant of 1000 kJ mol$^{-1}$ nm$^{-2}$ for production runs of 100 ns with both force field combinations. The simulation parameters recommended for CHARMM or Amber force fields in GROMACS were used [21]. Since these parameters are consistent between all simulations performed using the same force field, and they are listed below in a separate subsection.

The hydrogen bonding partners between the different TRAP subunits, between Sec61 and TRAP, between Sec61 and the ribosome, and between TRAP and the ribosome were calculated from the restrained simulations, in which the side chains were free to adapt to the environment. A hydrogen bond was defined by a donor–acceptor distance of 3.5 Å and a hydrogen–donor–acceptor angle of less than 30°. The calculations was performed using the HBonds tool in Visual Molecular Dynamics [17]. Key hydrogen bonds are listed in Tables S4, S5, and S7, and the key ones are highlighted in the structural snapshots in the main text. The last 90 ns were included in the analyses.

The simulations with a restrained backbone were also used to extract other key interactions between the lumenal domains of TRAP subunits. The `rerun` functionality of `gmx mdrun` was used to extract the short-range (no long-range electrostatics were included) Coulombic and van der Waals (Lennard-Jones potential) interactions. The major contributors to these energies are listed in Table S6.

##### All-atom simulations for protein dynamics and membrane–protein interactions

We studied the behavior of the complex formed by Sec61, TRAP, and the ribosome as well as its multiple subcomplexes. To this end, we embedded the atomistic models of 1) Sec61 alone, 2) TRAP alone, 3) Sec61 with TRAP, or 4) Sec61 with TRAP and parts of the ribosome into a lipid bilayer. Additionally, we performed a simulation of a protein-free bilayer as an additional control. The bilayer composition was set to mimic that of the ER membrane [1, 4, 6, 3], and it contained 54% phosphatidylcholine, 21% phosphatidylethanolamine, 10% phosphatidylinositol, 4% phosphatidylserine, 4% sphingomyelin, and 7% cholesterol. Since no information on lipid acyl chains was available, we modeled them as palmitate and oleate, except for sphingomyelin, which had a palmitate chain. For systems containing ribosome, the proteins and the parts of the RNA strands that were located in the vicinity of Sec61 or TRAP were included in the model. The sizes of the lipid membranes were adapted to the lateral extent of the protein, whereas the number of solvating waters was adjusted to solvate the entire protein. The system dimensions and molecule counts are provided in Table S1.

For the simulations containing parts of the ribosome, the ribosomal protein and RNA that lie far away from the Sec61 and TRAP were restrained to avoid modeling the entire ribosome, yet capturing the anchoring effect of its large size. The systems were generated in CHARMM-GUI [18, 34], downloaded in GROMACS-formats, and subjected to the standard equilibration protocol. We simulated all systems using the CHARMM family of force fields, namely CHARMM36m [16] for the protein, CHARMM36 for lipids [20, 33, 23] and RNA [8], and the CHARMM-compatible TIP3P model [19, 10] for water.

Additionally, the system containing all components (Sec61, TRAP, and parts of the ribosome) was also simulated using the Amber force fields, namely FF19SB [32] for the protein, Lipid21 [9] for the lipids, and the standard TIP3P model [19] for water. For the simulation with Amber force fields, not all lipid types found in the ER membrane were available in the lipid library, so for these simulation we adapted the composition to contain 62.75% phosphatidylcholine, 24.5% phosphatidylethanolamine, 4.75% phosphatidylserine, and 8% cholesterol.

The membrane systems with proteins were simulated for 2 µs each. The protein-free

| Model | # lipids | # water | dimensions ($x/y, z$) (Å) | atoms |
|---|---|---|---|---|
| **CHARMM force fields, membrane** | | | | |
| No protein | 1600 | 80000 | 213, 94 | 444834 |
| Sec61 | 800 | 80000 | 157, 140 | 350268 |
| TRAP | 800 | 80000 | 155, 144 | 353749 |
| Sec61+TRAP | 1000 | 100000 | 176, 140 | 447019 |
| Sec61+TRAP+ribos. | 1600 | 320000 | 220, 244 | 1205136 |
| **CHARMM force fields, bicelle** | | | | |
| Sec61+TRAP+ribos. | 1079 | 420000 | 252, 225 | 1445196 |
| **Amber force fields, membrane** | | | | |
| Sec61+TRAP+ribos. | 1600 | 320000 | 225, 236 | 1203354 |

Table S1: The summary of unrestrained atomistic MD simulations of the Sec61–TRAP–ribosome complex and its various subcomplexes. The membrane compositions between the CHARMM and Amber force field families differ somewhat depending on the availability of certiain lipid species in their lipid libraries (see text for detailed compositions). The dimensions are given in the membrane plane ($x/y$) or perpendicular to it ($z$)

membrane system was simulated for 500 ns. The simulation parameters were consistent within simulations using the same force field, and are listed below.

The protein stability was evaluated by calculating the root mean squared deviation (RMSD) of the protein backbone, after fitting the backbone structure first. This analysis was performed separately on Sec61 (all subunits together) or TRAP (all subunits together).

From these simulations involving full protein and lipid dynamics, we studied the membrane perturbations by the different protein assemblies. In long simulations, the membrane proteins rotate and their effects on the local membrane properties are smeared out. To account for this rotation, we first centered the protein, then RMSD-fitted the protein to a fixed orientation in the plane of the membrane. As such rotations would cause the membrane to cross the edges of the simulation box, the simulation box was simultaneously enlarged. We then performed the analyses with `g_lomepro` [12] on this larger system, and the last 1.5 µs were included in these spatial analyses. We extracted the leaflets shapes, the local membrane thickness, and the local order of the palmitate chains in each leaflet. Snapshots demonstrating the effect of the protein on the leaflet shape and membrane thickness were rendered using the tachyon rendered in Visual Molecular Dynamics [17]. Additionally, two-dimensional profiles of the leaflet position, membrane thickness, and acyl chain order were resolved by projecting the position, thickness, and order parameter maps onto a a line running parallel or perpendicular to the axis connecting Sec61 and TRAP. An area of 12 nm by 9 nm covered the entire extent of the protein, and was thus used for the projections.

Validation of the spatial analyses was performed by comparing the thickness and order parameter maps to those resolved from a protein-free system. The values in the map were histogrammed and fitted with a single (protein-free system) or a double (protein-containing systems) Gaussian.

We analyzed the openness of the lateral gate in Sec61 as a distance between the centers of mass of TM helices 2 and 7. This analysis was performed using `gmx distance` from the GROMACS simulation software [27].

##### All-atom simulations of bicelle curving

The complex formed by Sec61 and TRAP, and maintained in a certain conformation by the docked ribosome, seemed to induce local curvature in the membrane simulation. However, as the flat membrane cannot bend significantly due to the periodic simulation box, we repeated this simulation in a bicelle model. To this end, Sec61, TRAP, and parts of the ribosome were inserted in a POPC membrane and hydrated. We chose this single-component membrane to avoid any lipid demixing due to the bicelle edges. The system was generated in CHARMM-GUI [18, 34], and GROMACS-compatible simulation files were downloaded [21]. Then, we carved out a circularly shaped region with a diameter of ∼21 nm from the membrane, and subjected it to the standard CHARMM-GUI equilibration protocol. Then, the system was simulated for 1 µs using the suggested simulation parameters for CHARMM with GROMACS [21] with one exception: The compressibility of the barostat in the plane of the bicelle was set to 0 so the area in that plain was kept constant. The other used simulation parameters are listed below in detail. The bicelle system was simulated for 1 µs. The bicelle simulations were analyzed in the same manner as the membrane ones described above with g_lomepro [12].

##### Coarse-grained Simulations

Atomistic simulations are limited in size by the amount of computing power available. In a smaller membrane, significant curvature cannot build up due to the system periodicity. In the bicelle system, the bicelle will eventually drift away from the protein, limiting the achievable simulation time. Thus, we also generated a coarse-grained simulation model for the Sec61–TRAP complex. The coarse-grained protein was embedded in a lipid bilayer formed solely by POPC. This simple composition was chosen, as the recent Martini 3 force field [31] was used, and it currently lacks a vast and verified lipid library. The membrane contained a total of 3514 POPC molecules and it was solvated by ∼195,000 water beads. Neutralizing ions and ∼157 mM of NaCl were included. The system dimensions were $\sim 34 \times 34 \times 24$ nm$^3$. The system was simulated for 20 µs with the protein backbone restrained. These restraints locked the Sec61–TRAP complex into the conformation observed in the presence of the ribosome. The perturbations caused by the presence of the Sec61–TRAP complex were again evaluated using g_lomepro. Due to the restraints, no additional centering or alignment steps were necessary.

The ability of Sec61, TRAP, the different TRAP subunits, and the Sec61–TRAP complex to increase membrane permeability was probed by analyzing the phospholipid flip–flops. To this end, we also simulated Sec61 (all subunits included), TRAP (all subunits included), as well as the four TRAP subunits separately in a POPC membrane for 20 µs. In these simulations, the protein backbone structure was restrained. Details on all simulations are shown in Table S2. The flip–flops were analyzed based on the position of the lipid phosphate beads. A lipid was assigned to the upper (lower) leaflet, when this phosphate bead was at least 0.4 nm above (below) the membrane midplane. The coordinates were processed every

100 ns, and change in the leaflet identity was considered a flip–flop.

| Model | # lipids | # water | dimensions ($x/y, z$) | beads |
|---|---|---|---|---|
| Sec61 | 613 | 14316 | 146, 121 | 23276 |
| TRAP | 632 | 22142 | 146, 166 | 31909 |
| Sec61+TRAP | 3514 | 194652 | 234, 239 | 244146 |
| TRAP$\alpha$ | 655 | 21607 | 147, 160 | 30427 |
| TRAP$\beta$ | 654 | 17676 | 145, 141 | 26333 |
| TRAP$\gamma$ | 644 | 13523 | 145, 117 | 22041 |
| TRAP$\delta$ | 656 | 15727 | 148, 126 | 24342 |

Table S2: Summary of the coarse-grained simulations systems performed using the Martini 3 force field.

##### Simulation Parameters

The simulation parameters used with the different force fields are listed below in detail.

**All-atom CHARMM Family** The simulations were performed using the recommended simulation parameters for the CHARMM36 force field in GROMACS [21]. Namely, buffered Verlet lists were used to keep track of neighbour atoms [26]. The Lennard-Jones potential was cut off at 1.2 nm, and the forces were switched to zero between 1.0 nm and the cut-off distance. The smooth particle mesh Ewald (PME) algorithm was used for the calculation of long-range electrostatics [5, 11]. The temperature of the protein (and RNA), the membrane, and the solvent were separately maintained at 310 K using the stochastic velocity rescaling thermostat [2] with a time constant of 1 ps. The pressure along the membrane plane and normal to it were coupled to a semi-isotropic Parrinello–Rahman barostat [28] with a target pressure of 1 bar, compressibility of $4.5{\times}10^{-5}$ bar$-1$, and a time constant of 5 ps. The bonds involving hydrogen atoms were constrained using P-LINCS [14, 13], whereas water structure was constrained using SETTLE [24].

**All-atom Amber Family** The system was downloaded in GROMACS-compatible formats from CHARMM-GUI [21, 22], and subjected to the standard equilibration protocol of CHARMM-GUI. The system was simulated for 2 µs with an integration time step of 2 fs using GROMACS 2021 [27].

The simulation parameters provided by CHARMM-GUI were used [22]. Namely, buffered Verlet lists were used to track the neighbouring atoms for non-bonded interactions [26]. The Lennard-Jones potential was cut off at 0.9 nm, and a plain cutoff was used. Corrections due to the cutoff were performed to both energy and pressure [30]. The smooth particle mesh Ewald algorithm was used to calculate long-range electrostatics [5, 11]. The temperatures of the protein (with RNA), the lipids, and the solvent were maintained at 310 K by coupling them to a Nosé–Hoover thermostat [25, 15] with a time constant of 1 ps. The pressure was maintained at 1 bar with a semi-isotropic Parrinello–Rahman barostat [28] with a time constant of 5 ps and a compressibility of $4.5{\times}10^{-5}$ bar$^{-1}$. The bonds involving hydrogen atoms were constrained using P-LINCS [14, 13]. Waters were constrained with SETTLE [24].

**Coarse-grained Martini 3** The coarse-grained simulation systems were generated with the CHARMM-GUI Martini maker [29] and downloaded in GROMACS-compatible formats. The latest version 3.0 [31] of the Martini force field was used for the protein. All simulations were run for 20 µs with a time step of 20 fs using GROMACS 2021 [27].

We used the recently suggested “New-RF” simulation parameter set [7]. The Lennard-Jones potential was cut off at 1.1 nm, a distance at which the potential was shifted to zero. For electrostatics, a reaction field approach with a cutoff of 1.1 nm and a dielectric constant of $\infty$ was used for efficiency [7]. The stochastic velocity rescaling thermostat [2] with a time constant of 1 ps was applied separately to the protein, the lipids, and the solvent. A semi-isotropic Parrinello–Rahman barostat [28] with a time constant of 12 ps, compressibility of $3\times10^{-4}$ bar$^{-1}$, and a target pressure was applied semi-isotropically. Bonds were Electrostic interactions were screened by a dielectric constant of 15. Constraints present inherently in the force field were handled by P-LINCS [14, 13].

### S2 Supplementary Results

#### S2.1 Purification of the Sec61/TRAP translocon

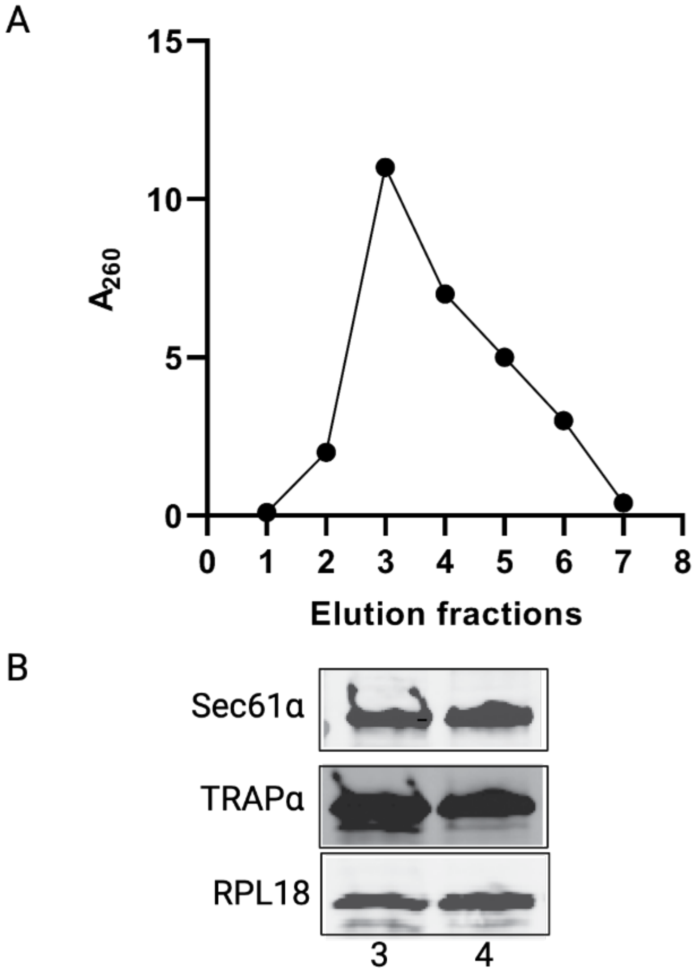


Figure S1.1: Purification of the Sec61/TRAP translocon bound to mammalian ribosome. **A)** Size exclusion chromatography of the solubilized ribosome/Sec61/TRAP complex with elution fractions from Superose-12 gel filtration assayed using absorbance at 260 nm. **B)** Western blot analysis of the elution fractions 3 and 4 using specific antibodies for Sec61$\alpha$, TRAP$\alpha$ and ribosomal rPL18.

#### S2.2 CryoEM data processing workflow

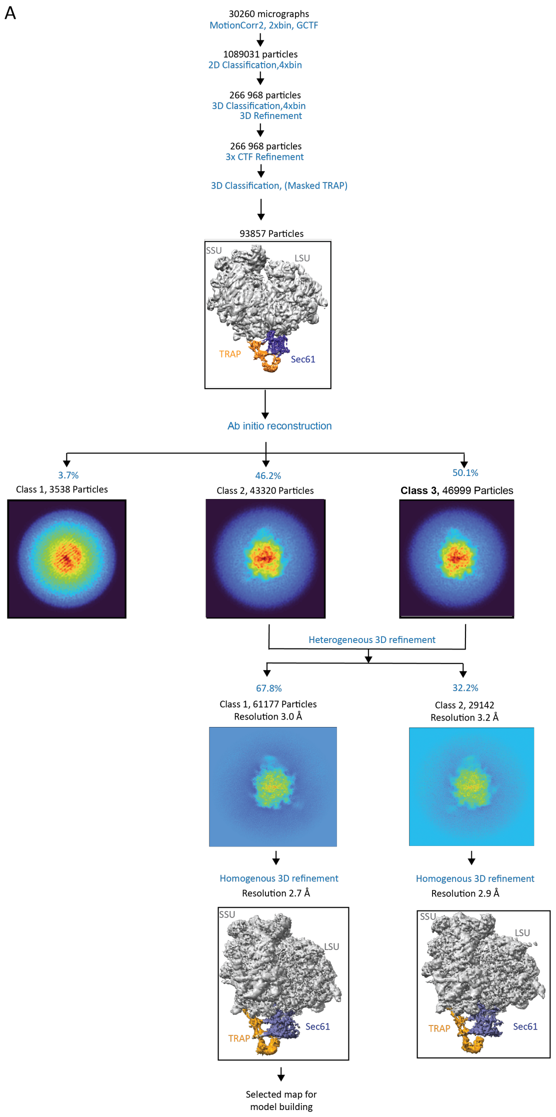


Figure S1.2: CryoEM data processing workflow. **A)** Schematic of pre-processing, classification and refinement procedures used to generate the Ribosome–Sec61–TRAP maps (see methods section for details). Maps shown after 3D classification and refinement processes highlight the Ribosome LSU and SSU in grey, the TRAP complex in orange and the Sec61 in blue. Image projection of ab initio reconstruction with three classes showing majority of the particles in class 2 and class 3. Image projections of heterogeneous refinement are shown and after further homogeneous refinement generated high resolution maps with the refined density of Ribosome-Sec61 and TRAP complex. *Ab initio* reconstruction, heterogeneous and homogeneous 3D refinement of the maps were processed in cryoSPARC, and the rest of the jobs were processed in Relion 3.046 maintained within the Scipion v3.0.7.

#### S2.3 Local resolution

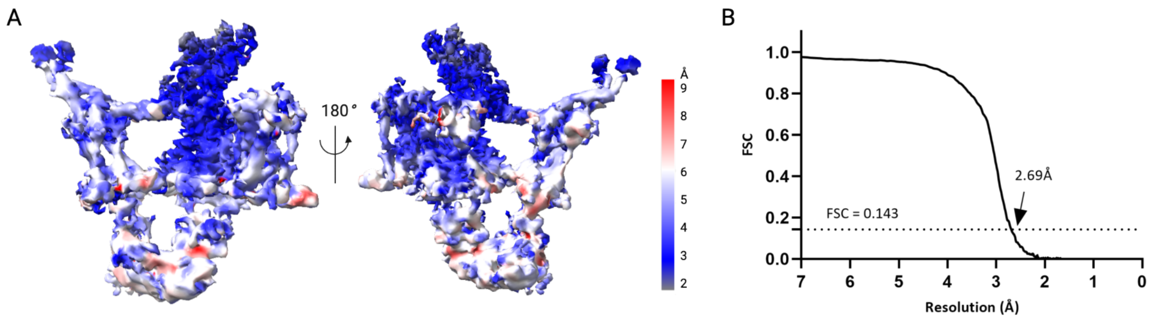


Figure S1.3: FSC curve and the estimation of local resolution **A)** Density maps of the structure of Ribosome-Sec61-TRAP coloured by local resolution estimation with cryoSPARC and ChimeraX. **B)** FSC curve as a function of resolution using output from homogeneous refinement in cryoSPARC v3.3.2.

#### S2.4 AlphaFold models

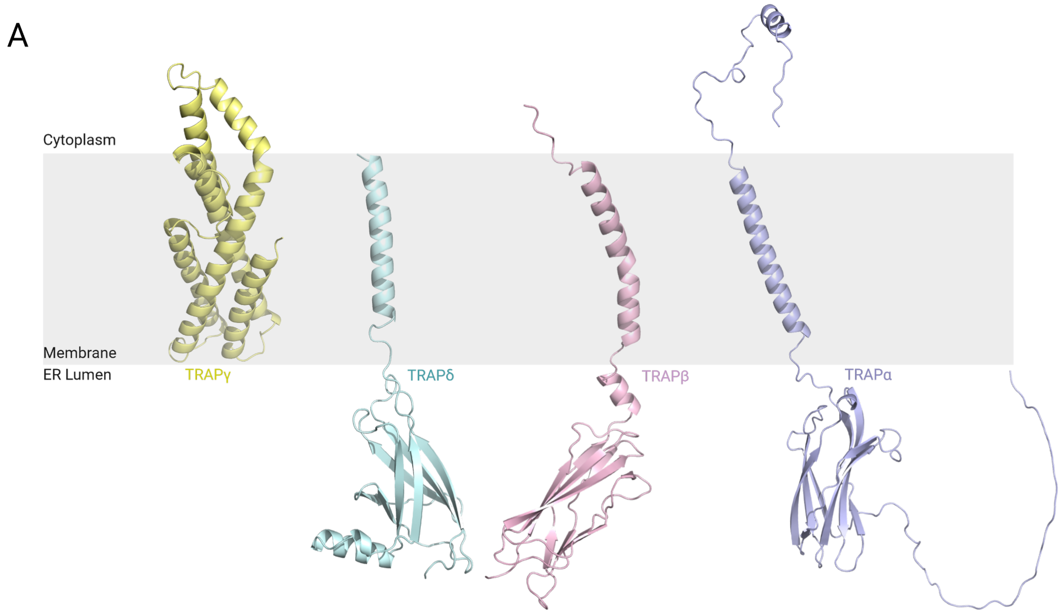


Figure S1.4: AlphaFold2 models of the TRAP subunits. **A)** AlphaFold2 model of TRAP$\gamma$ (yellow) with a bundle of 6 helices in the membrane and cytoplasmic region, TRAP$\delta$ (cyan) with small folded beta sheet rich domain in the ER lumen followed by a short linker to the single helix in the ER membrane, TRAP$\beta$ (pink) contains a small folded lumenal domain similar to TRAP$\delta$ in ER lumen, followed by a single TM domain and short tail in the cytoplasmic region, and TRAP$\alpha$ (blue) has an unstructured region in the N-terminal followed by a small folded domain in the lumenal region, includes single transmembrane domain followed by cytoplasmic region that is mostly unstructured in the model.

##### S2.5 Fit of the individual TRAP subunit models in the cryo-EM density

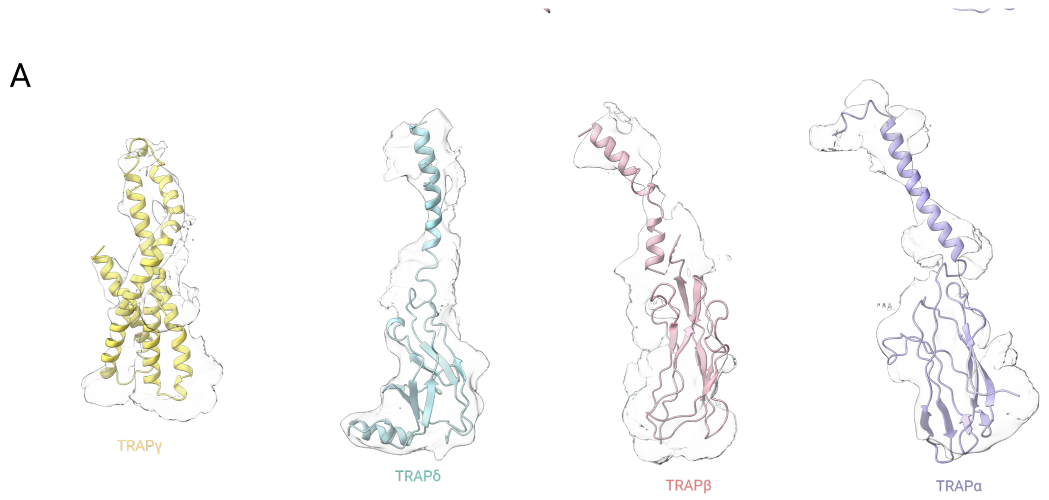


Figure S1.5: TRAP subunits structures into the cryoEM density map. **(A)** Representative cryo-EM density fragments with the final refined TRAP subunit models.Density map shown is locally filtered homogenous refinement output map from cryoSPARC (sigma value : 0.6).

#### S2.6 Density of the N-terminal TRAP$\alpha$

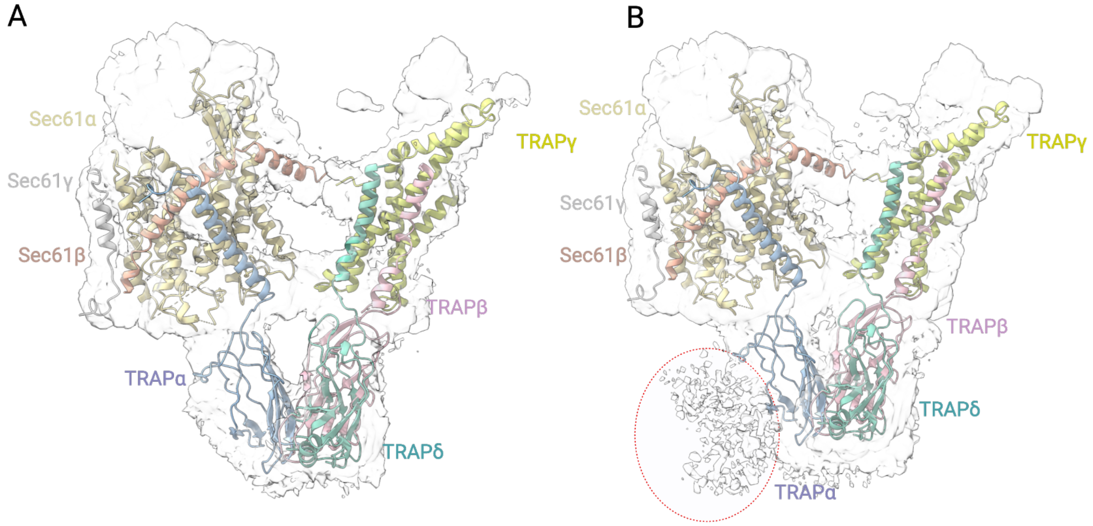


Figure S1.6: Cryo-EM density of the TRAP$\alpha$ N-terminus. **A)** Cryo-EM map of the TRAP complex obtained before 3D focused classification (detail Fig S1.2) with Sec61/TRAP structure, sigma value: 0.2. **B)** At a higher contour level (sigma value 1), an extra weak density representing the unstructured N-terminus of TRAP$\alpha$ can be visualized

##### S2.7 Fit of the Sec61/TRAP atomic model to the cryo-ET density

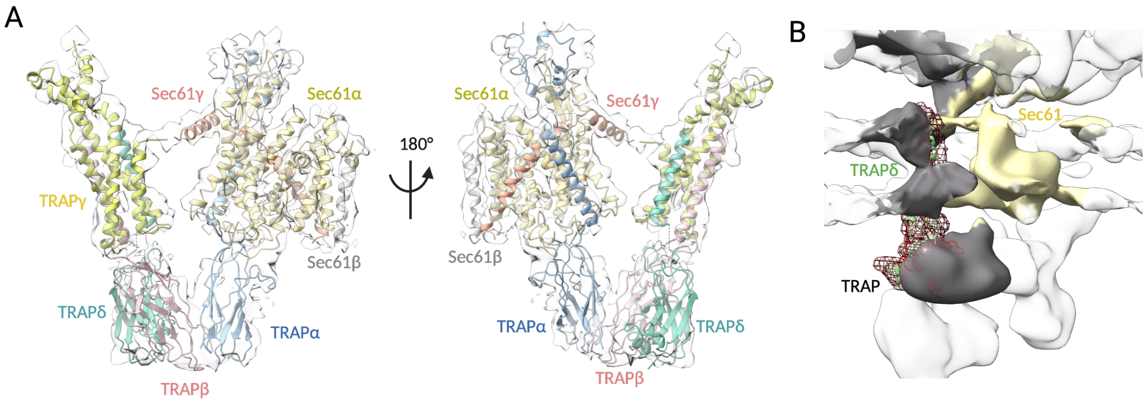


Figure S1.7: Fit of the Sec61/TRAP atomic model to the cryo-ET density of Sec61/TRAP in the ER membrane. **A)** Cryo-ET density (EMD-3068) of TRAP/Sec61 from subtomogram averaging in intact ER membranes and the fit of our single particle cryo-EM model fitted onto the cryo-ET density. TRAP subunits are colored as TRAP$\alpha$:cyan, TRAP$\beta$:pink, TRAP$\delta$:green, and TRAP$\gamma$: yellow. **B)** Difference density originating from TRAP$\delta$-deficient fibroblast translocon EMD-4143 (surface representation, TRAP: grey and Sec61: yellow) and the isolated density of the TRAP complex from EMD-3068 (red mesh).

#### S2.8 Glycosylation sites on TRAP$\alpha$ and TRAP$\beta$

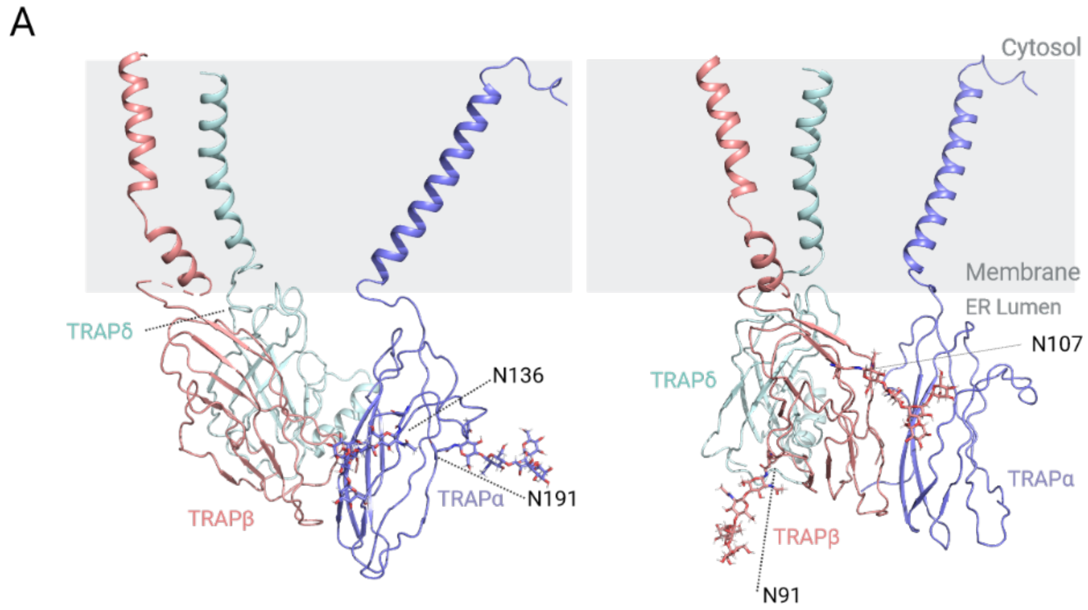


Figure S2: Glycosylation sites on TRAP$\alpha$ and TRAP$\beta$. **A)** Cartoon representation of the TRAP$\alpha$ and TRAP$\beta$ with predicted glycans indicated. Figure in the left shows the TRAP complex with the two glycosylation sites of TRAP$\beta$, glycans are color-coded as carbon: light red, oxygen: red. In the right shows the TRAP complex with the two glycosylation sites of TRAP$\alpha$, glycans are color-coded as carbon: blue, oxygen: red.

##### S2.9 Refinement and model statistics

| RNC-Sec61-TRAP | |
|---|---|
| **Data Collection and Processing** | |
| Magnification | 105000 |
| Voltage | 300 |
| Electron exposure ($e^{-1}/Å^2$) | 47.66 |
| Defocus range (µm) | $-0.6$ to $-2.2$ |
| Pixel size (Å) | 0.415 (super resolution) |
| Symmetry imposed | C1 |
| Initial particles | 1089031 |
| Final particles | 61177 |
| Map resolution (Å) | 2.69 |
| Fourier Shell Correlation threshold | 0.143 |
| **Refinement** | |
| Map pixel size (Å) | 0.83 |
| Model Resolution (Å) | 2.69 |
| **Model Composition** | |
| Atoms | 64453 (Hydrogens: 255) |
| Protein/Nucleotide residues | 2122/2200 |
| Ligands | NME |
| | ACE |
| **Root-Mean-Square Deviations (RMSDs)** | |
| Bond lengths (Å) | 0.006 |
| Bond angles ($^{c}irc$) | 0.712 |
| **Validation** | |
| MolProbity score | 2.68 |
| Clashscore | 26.02 |
| **Ramachandran Plot** | |
| Favoured (%) | 92.82 |
| Allowed (%) | 7.18 |
| Disallowed (%) | 0.00 |

Table S3: Refinement and Model Statistics

##### S2.10 Supplementary results from MD simulations

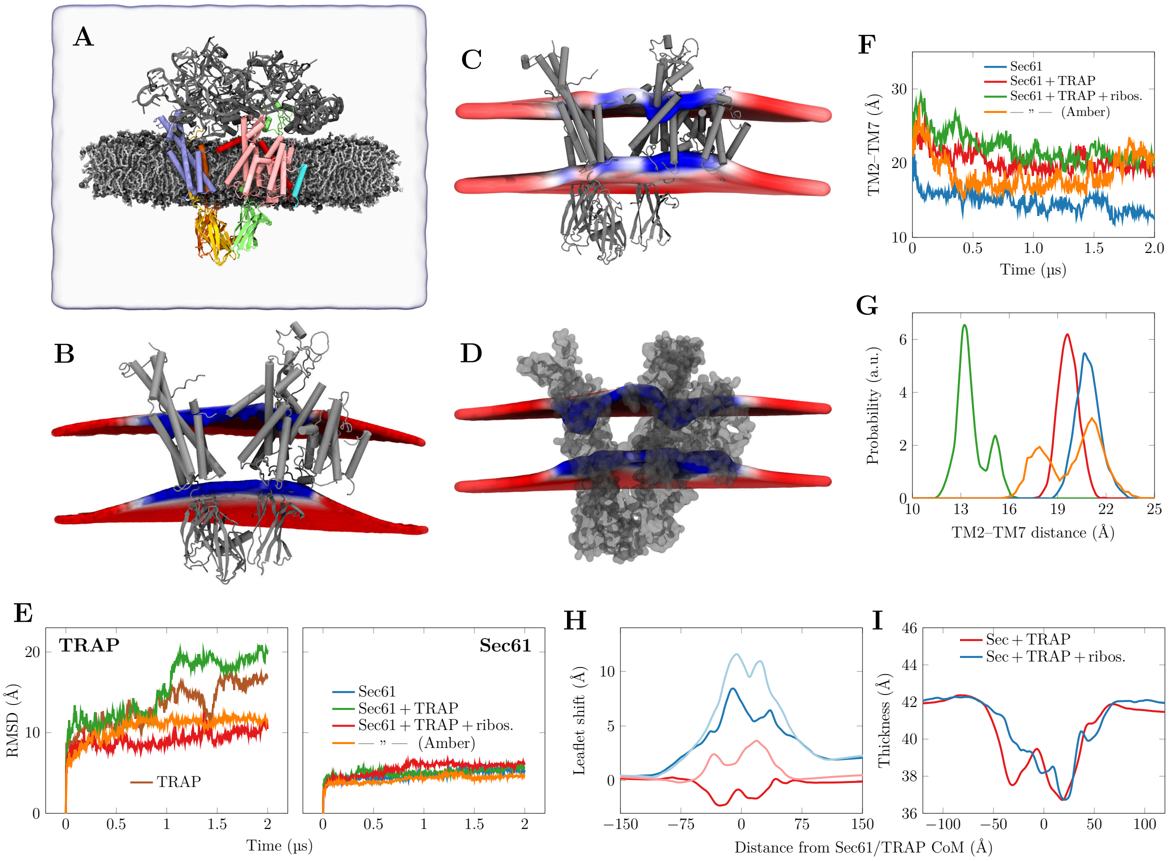


Figure S3: **A)** Snapshot of the bicelle simulation system after equilibration protocol. The membrane consists purely of POPC, which is shown in light gray licorice with the phosphorus atoms drawn in dark gray spheres. The water and thus the simulation box extent is shown as a transparent blue surface. The ribosomal proteins and RNA re coloured in dark gray. TRAP$\alpha$ is drawn in green, TRAP$\beta$ in yellow, TRAP$\gamma$ in blue, TRAP$\delta$ in orange, Sec61$\alpha$ in pink, Sec61$\beta$ in cyan, and Sec61$\gamma$ in dark red. Lipid hydrogens and ions are omitted for clarity. **B)** Membrane lensing of the Sec61/TRAP complex in the presence of the ribosome in the atomistic POPC bicelle. The phoshorus locations of the two leaflets are shown by the surfaces, and the colour shows local thickness (red: 43 Å, blue: 37 Å). The lower (lumenal) leaflet shows significant curvature. **C)** The membrane lensing in the atomistic simulation of Sec61/TRAP complex in the presence of the ribosome in an ER membrane with the Amber force field family. The result is similar to that obtained with CHARMM36 force fields (main text). Coloring from XX (blue) to YY Å (red). **D)** The membrane lensing in the simulation of the backbone-restrained Sec61/TRAP complex simulated in the coarse-grained scheme using the Martini 3 force field. The result is similar to that obtained with atomistic force fields. Colouring from 34 (blue) to 40 Å (red). **E)** The time evolution of RMSD values of TRAP and Sec61. Unlike the corresponding figure in the main text, this on contains data for the Sec61/TRAP system in the presence of the ribosome and simulated with the Amber force fields. **(Continues on the next page)**

Figure S3: **F)** Lateral gate openness characterized by the distance of the centers of mass of TM helices 2 and 7. Data for the Sec61/TRAP system in the presence of the ribosome and simulated with the Amber force fields is also included here. **G)** Histogram of the distance in F) extracted during the last 500 ns of the simulations. **H)** The positioning of the leaflets in the systems with the Sec61/TRAP in the presence (blue) or absence (red) of the ribosome. The darker (lighter) lines show data for the cytosolic (lumenal) leaflet. Curvature is only induced when ribosome anchors TRAP in the specific V-shaped conformation. The data is collected from an elongated membrane patch covering the extent of the Sec61/TRAP TM domains, and aligned parallel to the axis connecting Sec61 and TRAP. **I)** Local membrane thickness. Although the Sec61/TRAP system without the ribosome is not curved, the thinning caused by the protein hydrophobic mismatch is still similar for the systems with and without ribosomal anchoring.

#### S2.11 Hydrogen bonding between Sec61 and TRAP

| Donor | Acceptor | CHARMM36m | FF19SB | Comments |
|---|---|---|---|---|
| | | **Sec61$\alpha$–TRAP$\alpha$** | | |
| TYR235-Side | GLU204-Side | 93% | 86% | |
| ARG205-Side | GLU168-Side | 69% | 66% | |
| | | **Sec61$\alpha$–TRAP$\gamma$** | | |
| LYS158-Side | ASP357-Side | 33% | 79% | |
| | | **Sec61$\gamma$–TRAP$\gamma$** | | |
| PHE7-Main | LYS185-Side | 34% | 54% | N-terminal PHE7 & |
| PHE7-Main | LYS185-Main | 32% | 4% | C-terminal LYS185 |
| VAL8-Main | LYS185-Side | 42% | 59% | C-terminal LYS185 |
| VAL8-Main | LYS185-Main | 22% | 0% | C-terminal LYS185 |

Table S4: Key hydrogen bonds between Sec61 and TRAP in the backbone-restrained simulations using two complementary sets of protein and lipid force fields. The occupancies observed with both force fields are also listed. Notably, TRAP$\gamma$ and Sec61$\alpha$ are at a distance where hydrogen bonding is possible, but no hydrogen bonds meet the above criteria.

##### S2.12 Hydrogen bonding between TRAP subunits

| Donor | Acceptor | CHARMM36m | FF19SB | Comments |
|---|---|---|---|---|
| | | **TRAP$\alpha$–TRAP$\beta$** | | |
| ARG150-Side | GLU119-Side | 28% | 62% | Lumenal domains |
| ARG150-Side | GLU119-Main | 9% | 14% | Lumenal domains |
| | | **TRAP$\gamma$–TRAP$\delta$** | | |
| LYS91-Side | ALA173-Main | 46% | 59% | Terminal ALA173 |
| LYS91-Side | ALA173-Side | 34% | 12% | Cytosolic membrane interface |
| | | **TRAP$\beta$–TRAP$\gamma$** | | |
| ARG58-Side | GLU137-Side | 89% | 84% | Lumenal membrane interface |
| SER167-Side | ASN142-Side | 32% | 86% | Membrane core |
| | | **TRAP$\beta$–TRAP$\delta$** | | |
| ILE53-Main | ASP82-Side | 86% | 87% | Lumenal domains |
| ARG39-Side | GLU150-Side | 83% | 46% | Lumenal membrane interface |
| ASN52-Side | ASP82-Side | 73% | 85% | Lumenal domains |
| SER35-Side | GLU46-Side | 52% | 64% | Lumenal domains |
| SER47-Side | LEU29-Main | 0% | 81% | Lumenal domains |

Table S5: Key hydrogen bonds between TRAP subunits in the backbone-restrained simulations using two complementary sets of protein and lipid force fields. The occupancies observed with both force fields are also listed.

#### S2.13 Key interactions between TRAP subunits

| Residue | CHARMM36m | FF19SB |
|---|---|---|
| **TRAP**$\alpha$ **with TRAP**$\beta$, total of $-338.0$ / $-326.6$ kJ/mol | | |
| SER82 | -83.4 | -81.4 |
| ARG150 | -67.4 | -77.5 |
| **TRAP**$\beta$ **with TRAP**$\alpha$, total of $-338.0$ / $-326.6$ kJ/mol | | |
| GLU21 | -78.7 | -84.0 |
| GLU119 | -69.8 | -80.0 |
| **TRAP**$\beta$ **with TRAP**$\delta$, total $-614.2$ / $-642.9$ kJ/mol | | |
| ASN48 | -81.0 | -71.0 |
| ASN34 | -58.8 | -59.1 |
| PRO88 | -55.9 | -57.5 |
| **TRAP**$\delta$ **with TRAP**$\beta$, total $-614.2$ / $-642.9$ kJ/mol | | |
| ASP82 | -94.9 | -88.8 |
| ARG79 | -82.0 | -98.2 |
| GLU46 | -43.3 | -43.4 |

Table S6: The most dominating residues involved in the interaction between the lumenal domains of TRAP subunits. The data are extracted from backbone-restrained simulations using two complementary sets of protein and lipid force fields. Only residues that interact with more than 41.84 kJ/mol (10 kcal/mol) in both simulation force fields are listed. The total values (given as CHARMM/Amber) are calculated over interacting residues, not only the ones listed here.

#### S2.14 Hydrogen bonding between Sec61/TRAP and the ribosome

| Donor | Acceptor | CHARMM36m | FF19SB | Comments |
|---|---|---|---|---|
| | **TRAP$\alpha$–RNA. 8** | | | |
| LYS235-Side | A84-Side | 66% | 61% | |
| | **TRAP$\gamma$–RNA. 5** | | | |
| ARG114-Side | G2550-Side | 73% | 60% | |
| ARG110-Side | U2763-Side | 59% | 56% | |
| | **Sec61$\alpha$–RNA 5** | | | |
| ARG405-Side | C2526-Side | 85% | 76% | |
| ARG273-Side | G2433-Side | 53% | 48% | |
| LYS268-Side | U2432-Side | 39% | 50% | |
| | **Sec61$\gamma$–Ribosomal L23a** | | | |
| LYS16-Side | ASP148-Side | 76% | 17% | |
| ARG20-Side | ASP148-Side | 93% | 82% | |
| ARG24-Side | ILE156-Main | 88% | 5% | C-terminal ILE156 |
| ARG24-Side | ILE156-Side | 86% | 57% | C-terminal ILE156 |
| ASN151-Side | ARG20-Main | 62% | 65% | |
| | **Sec61$\alpha$–Ribosomal L23a** | | | |
| SER408-Side | GLU84-Side | 90% | 37% | |
| GLU84-Main | GLY403-Main | 62% | 72% | |
| TYR416-Side | ILE156-Side | 56% | 95% | C-terminal ILE156 |
| | **TRAP$\gamma$–Ribosomal L38** | | | |
| ARG16-Side | GLU119-Side | 16% | 84% | |

Table S7: Hydrogen bonds between Sec61 and the ribosomal proteins and RNA chains in the backbone-restrained simulations using two complementary sets of protein and lipid force fields. The occupancies observed with both force fields are also listed. Notably, ribosomal proteins L35, L39, and L19 lie in the vicinity of Sec61$\alpha$, yet none of the observed hydrogen bonds meet the above criteria. The same holds true for the TRAP$\gamma$ and ribosomal protein L38, as well as Sec61$\gamma$ and ribosomal protein L35.
